# *SMARCA4* mutation induces tumor cell-intrinsic defects in enhancer landscape and resistance to immunotherapy

**DOI:** 10.1101/2024.06.18.599431

**Authors:** Yawen Wang, Ismail M Meraz, Md Qudratullah, Sasikumar Kotagiri, Yanyan Han, Yuanxin Xi, Jing Wang, Yonathan Lissanu

## Abstract

Cancer genomic studies have identified frequent alterations in components of the SWI/SNF (SWItch/Sucrose Non- Fermenting) chromatin remodeling complex including *SMARCA4* and *ARID1A*. Importantly, clinical reports indicate that *SMARCA4*-mutant lung cancers respond poorly to immunotherapy and have dismal prognosis. However, the mechanistic basis of immunotherapy resistance is unknown. Here, we corroborated the clinical findings by using immune-humanized, syngeneic, and genetically engineered mouse models of lung cancer harboring *SMARCA4* deficiency. Specifically, we show that *SMARCA4* loss caused decreased response to anti-PD1 immunotherapy associated with significantly reduced infiltration of dendritic cells (DCs) and CD4+ T cells into the tumor microenvironment (TME). Mechanistically, we show that *SMARCA4* loss in tumor cells led to profound downregulation of *STING, IL1β* and other components of the innate immune system as well as inflammatory cytokines that are required for efficient recruitment and activity of immune cells. We establish that this deregulation of gene expression is caused by cancer cell-intrinsic reprogramming of the enhancer landscape with marked loss of chromatin accessibility at enhancers of genes involved in innate immune response such as *STING, IL1β,* type I IFN and inflammatory cytokines. Interestingly, we observed that transcription factor NF-κB binding motif was highly enriched in enhancers that lose accessibility upon *SMARCA4* deficiency. Finally, we confirmed that SMARCA4 and NF-κB co-occupy the same genomic loci on enhancers associated with *STING* and *IL1β,* indicating a functional interplay between SMARCA4 and NF-κB. Taken together, our findings provide the mechanistic basis for the poor response of *SMARCA4*-mutant tumors to anti-PD1 immunotherapy and establish a functional link between SMARCA4 and NF-κB on innate immune and inflammatory gene expression regulation.

## Introduction

Lung cancer is a devastating disease that remains the top cause of cancer mortality (1, 2). Despite the advent of targeted therapy against oncogenic kinases and recent KRAS^G12C^ inhibitors, the majority of patients with lung cancer still lack effective therapeutics, underscoring the dire need for additional treatment options (3, 4). Significant strides have been made in recent years with the development of immune checkpoint inhibitors targeting CTLA4 and PD-1/PD-L1 improving the outcomes for some lung cancer patients (5, 6).

The SWI/SNF (SWItch/Sucrose Non- Fermenting) chromatin remodeling complex is a large multi-protein assembly that uses the energy derived from ATP hydrolysis to remodel nucleosomes and facilitate chromatin dependent cellular processes such as DNA replication, repair and transcription (7–9). Subunits of the SWI/SNF chromatin remodeling complex including *SMARCA4* and *ARID1A* are frequently mutated in lung cancer (16% in early-stage disease as in TCGA and up to 33% in advanced stages) (10–13). Several recent independent clinical studies have shown that *SMARCA4* mutant lung cancers have one of the worst prognosis among genetically defined subtypes of lung cancer (6, 14–16). Moreover, *SMARCA4* mutant tumors have poor response to immunotherapy (6, 14). Bioinformatic analysis into the tumor microenvironment of *SMARCA4* mutant lung cancer revealed these tumors have significantly lower levels of infiltrating cytotoxic T-cells and show “immune cold” features (6, 14) which is in line with the lack of clinical response to immunotherapy. In addition to primary resistance to immunotherapy, *SMARCA4* mutation was recently identified as the second most common genetic alteration upon development of acquired resistance to immune checkpoint inhibitors in lung cancer(17), strongly suggesting a key role of *SMARCA4* in modifying anti-tumor immunity. Importantly, *SMARCA4* mutant tumors have high tumor mutational burden (TMB) (18, 19), thus the lack of immunotherapy response is particularly puzzling. To date, the molecular mechanisms of resistance of *SMARCA4* mutant tumors to immunotherapy remain unknown.

The most commonly clinically utilized immunotherapies are T cell-centered (20–25). However, the effector functions of T cells are non-autonomous and studies have shown that the initiation and sustainability of T cell response and the maintenance of T cell memory depend on interactions with other immune cells such as dendritic cells (DCs) and innate immunity (26–30). Importantly, a growing body of work has recently shown critical role of DCs in anti-tumor immunity and shaping T cell activity during immunotherapy (31–36). In these processes, interferons (IFNs) and inducers of IFN response such as STING play a key role(32, 37). Specifically, innate immune sensing of tumors largely occurs through the host STING pathway, which leads to type I interferon (IFN) production, dendritic cell (DC) activation, cross-presentation of tumor-associated antigens to CD8+ T cells and T cell recruitment into the tumor microenvironment (38, 39). This has generated tremendous excitement and efforts to re-engage the innate immune system using STING agonists are underway (40–43). Furthermore, therapies aimed at activating DCs such as by administration of Fms-Like tyrosine kinase 3 (Flt3) ligand, a growth factor promoting DCs development, in combination with αCD40 (to activate DCs), termed DC therapy, are being actively investigated (31, 44).

Here, we show that *SMARCA4* mutant tumors respond poorly to anti-PD1 immunotherapy by using several orthogonal models of lung cancer. We show that *SMARCA4* loss leads to significantly reduced infiltration of DCs and CD4+ T cells into the tumor microenvironment. Mechanistically, we show that loss of *SMARCA4* in cancer cells results in a profound epigenetic deregulation of enhancers leading to downregulation of *STING, IL1β* and other components of the innate immune system that are required for efficient recruitment and activity of immune cells. We also show that SMARCA4 and NF- κB co-occupy the same genomic loci on enhancers associated with *STING* and *IL1β,* indicating a functional interplay between SMARCA4 and NF-κB. Taken together, our findings provide a cancer cell-intrinsic mechanistic basis for the poor response of *SMARCA4*-mutant tumors to anti-PD1 immunotherapy and establish a functional link between SMARCA4 and NF-κB on innate immune and inflammatory gene expression regulation.

## Results

### *SMARCA4* deficiency leads to decreased response to anti-PD1 immunotherapy and impaired immune cell infiltration into tumors

To study the role of *SMARCA4* loss on anti-tumor immunity, we generated immune-humanized mouse models by reconstitution of umbilical cord derived purified CD34+ hematopoietic stem cells (HSC) into irradiated NSG mice (termed hu-NSG from here on) as described previously (45). Briefly, freshly isolated CD34+ HSCs from cord blood were injected intravenously into mice after 24 hours irradiation (Fig. 1A). After 4-6 weeks, mice that had over 25% human CD45+ cells in the peripheral blood were considered humanized, hu-NSG mice. Moreover, human immune cell populations including CD3+ T cells, CD19+ B cells, CD56+ NK cells were determined in the hu-NSG mice (Supplementary Fig. S1).

**Figure 1:**
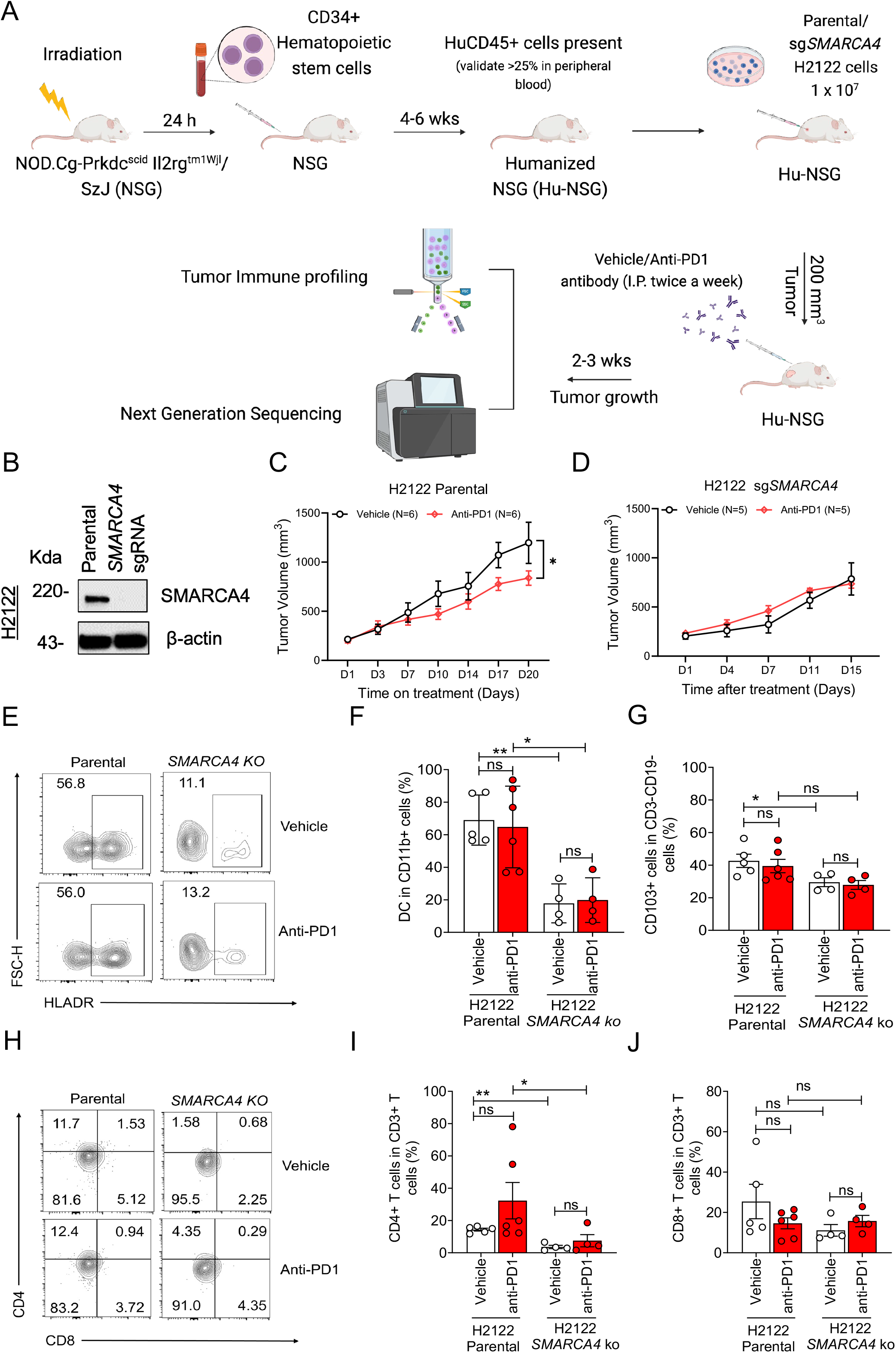
*SMARCA4* deficiency leads to decreased response to anti-PD1 immunotherapy and impaired immune cell infiltration into tumors in humanized xenograft models. (**A**) Schematic representation detailing the development of the humanized mouse model. Parental and *SMARCA4* KO H2122 tumors were introduced into humanized mice at 6 weeks post-implantation, with a minimum of 25% huCD45+ cells observed in peripheral blood. Parental xenograft models were subjected to i.p. injections of vehicle or pembrolizumab (250 µg/mouse) every 3 days for 3 weeks, while *SMARCA4* KO xenograft models were treated in the same manner but for 2 weeks. (**B**) Immunoblots of parental and *SMARCA4* KO H2122 cell lysates stained using anti-SMARCA4 and β-actin antibodies. Protein molecular weight markers are in kilodaltons (kDa). **(C-D)** Growth curves of parental H2122 xenograft models (n = 6 animals per group) (**C**) or *SMARCA4* KO H2122 xenograft models (n = 5 animals per group) (**D**) post-treatment with isotype control antibodies or anti-PD1 antibodies were generated. Data points represent mean ± S.E.M., * *p* <0.05, two-sided unpaired t-test. (**E-J**) The cells isolated from dissociated subcutaneous parental and *SMARCA4* KO H2122 xenograft models were analyzed by flow cytometry. **(E-G)** Flow cytometry plot of HLA-DR antibody **(E).** The percentage of Dendritic cells (DCs) out of CD11b+ cells **(F)** and CD103+ cells out of CD3-CD19-cells **(G)**. **(H-J)** Flow cytometry plot of CD4 and CD8 antibodies **(H).** The percentage of CD4+ T cells **(I)** and CD8+ T cells **(J)** out of CD3+ T cells. Data points represent mean ± S.E.M., \*\**p*<0.01, \**p* <0.05, One-tailed Mann-Whitney test.

We next injected isogenic pair of parental and *SMARCA4* knockout H2122 human lung cancer cells subcutaneously into hu-NSG mice (Fig. 1A-B). H2122 cells have been reported to exhibit moderate response to anti-PD1 treatment (46). When tumors reached 200mm^3^, hu-NSG mice were treated with vehicle or anti-PD1 antibodies every 3 day for 2-3 weeks. While parental tumors showed moderate but significant tumor growth inhibition, *SMARCA4* knockout tumors were fully resistant to anti-PD1 (Fig. 1C-D).

Next, we determined the repertoire of tumor infiltrating immune cells by using flow cytometry-based immune profiling of dissociated tumors. This revealed a significant reduction of total dendritic cells (DCs), CD103+ cDC1 cells, CD4+ T-cells and trend towards reduced CD8+ T cells in *SMARCA4* knockout tumors (Fig. 1 E-J). Additionally, *SMARCA4* deficiency significantly reduced percentage of CD3+ T cells, CD4+CD8+ T cells, Treg cells, PD-1 NK cells, myeloid-derived suppressor cells (MDSC), CD11b+ cells and M2 macrophages, but no obvious effect on other T cell subtypes, B cells, NK cells, monocytes, or M1 macrophages (Supplementary Fig. S2).

To confirm and expand on these observations, we utilized an orthogonal experimental model. Specifically, we knocked out *Smarca4* in FM471 (47) mouse lung cancer cells (Fig. 2B). These cells were derived from lung adenocarcinomas of C57/BL6 mice subjected to tobacco smoke, have *Kras* and *p53* mutations and harbor a high tumor mutational burden (TMB) and thus are excellent model system to investigate the immune system in immunocompetent mice against a high TMB tumor type such as lung cancer. We injected parental and isogenic *Smarca4* knockout FM471 cells subcutaneously in C57/BL6 mice and treated mice with isotype control or anti-PD1 antibodies for 4 doses (Fig. 2A). While parental tumors showed robust response to anti-PD1 with greater than 50% tumor growth inhibition, *Smarca4* knockout tumors had minimal response of about 20% reduction (Fig. 2C and Supplementary Fig. S3A). We then performed comprehensive FACS profiling of infiltrating immune cells. Similar to our previous observation, the most consistent and significant differences were profound reduction of total dendritic cells (DCs), cDC1 cells (Fig. 2D-F and Supplementary Fig. S3B) and CD4+ T-cells in *Smarca4* knockout tumors (Fig. 2G-I). In addition, *Smarca4* deficiency moderately reduced KLRG1+CXCR3-TEFF cells, CD8+PD1+ T cells, CD8+CD69+ T cells in TME (Supplementary Fig. S3-5).

**Figure 2:**
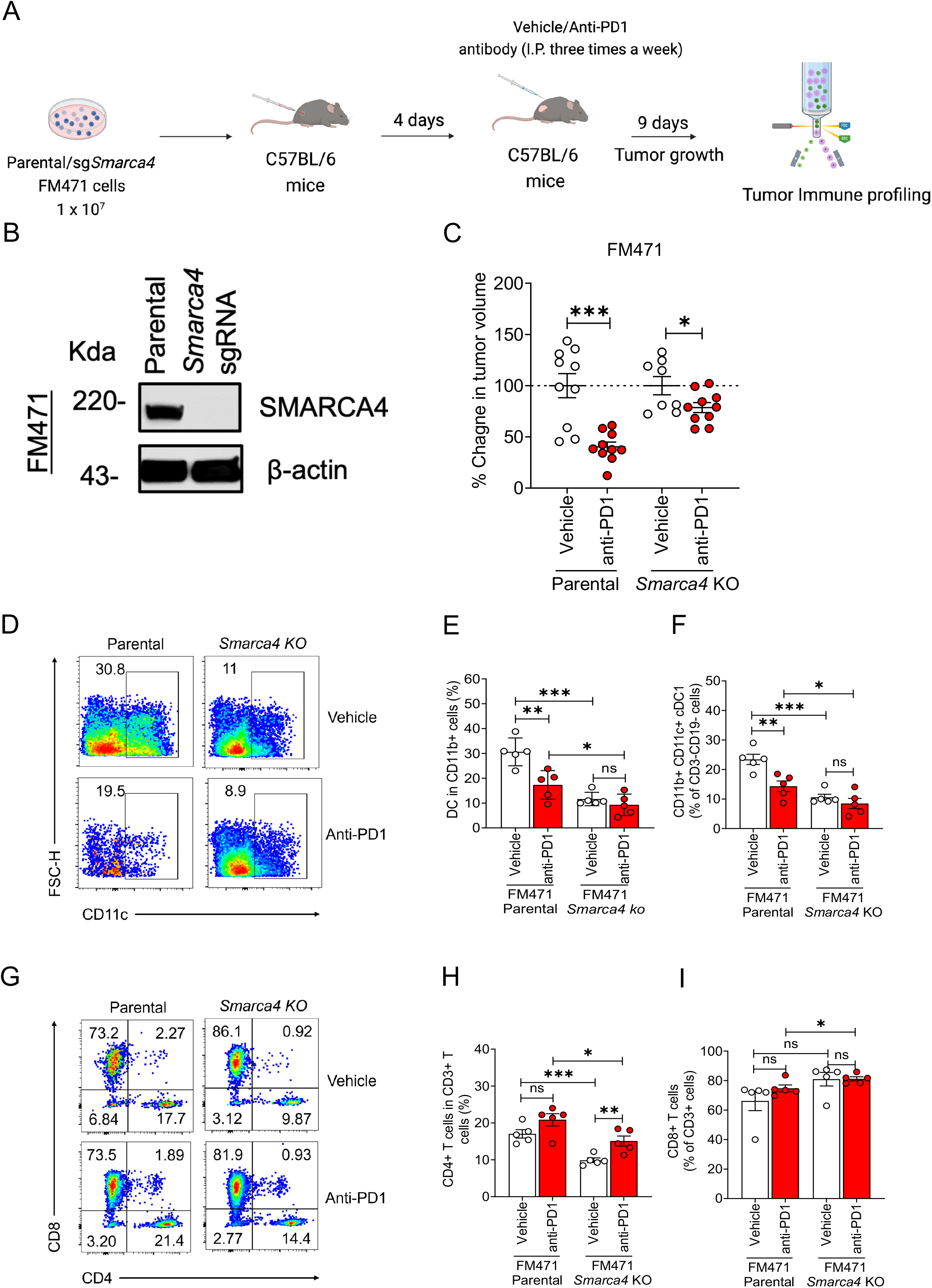
*Smarca4* deficiency leads to decreased response to anti-PD1 immunotherapy and impaired immune cell infiltration into tumors in mouse syngeneic models. (**A**) Schematics of immunotherapy experiments setup for FM471 tumor model. Parental and *Smarca4* KO FM471 cancer cell lines were subcutaneously injected into eight-week-old C57BL/6 female mice (n = 20 animals per group). 9 days isotype control antibodies or anti-PD1 antibodies were treated after tumors were established at day 4 after implantation. (**B**) Immunoblots of parental and *Smarca4* KO FM471 cell lysates stained using anti-SMARCA4 and β-actin antibodies. Protein molecular weight markers are in kilodaltons (kDa). (**C**) Analysis of the percent change in tumor volume of parental and *Smarca4* KO FM471 syngeneic models post-treatment with isotype control antibodies or anti-PD1 antibodies. (**D-I**) Cells were isolated from dissociated subcutaneous parental and *Smarca4* KO FM471 syngeneic models. **(D-F)** Flow cytometry plot of total CD11c+ cells **(D)**. Percentage of (**E**) DCs defined by total CD11c+ gating from CD11b+ cells and (**F**) cDC1 cells define by inter medium CD11c+ and CD11b+ cells. **(G-I)** Flow cytometry plot of CD4 and CD8 (**G**). The percentage of CD4+ T cells **(H)** and CD8+ T cells **(I)** out of CD3+ T cells. Data points represent mean ± S.E.M., \*\*\**p* < 0.001, \*\**p*<0.01, \**p* <0.05, two-sided unpaired t-test.

Taken together, the two orthogonal model systems showed consistent and significant reduction in DCs and CD4+ T cells while frequency of many other immune cells were not changed, minimally altered or showed mixed alteration. Hence, we decided to focus our next investigations on how loss of *SMARCA4* in cancer cells results in diminished DCs and CD4+ T cells within the TME.

To gain deeper insights into the mechanistic basis for the impaired response to immunotherapy and compromised recruitment of DCs and T-cells, we performed RNA-seq on H2122 parental and *SMARCA4* knockout tumors. Differential gene expression analysis showed 485 genes were upregulated and 612 were downregulated in *SMARCA4* knockouts with *SMARCA4*, as expected, being the most downregulated gene (Fig. 3A-B). Unbiased gene set enrichment analysis (GSEA) of the most significantly downregulated genes in *SMARCA4* knockout tumors showed interferon alpha (IFNα) and interferon gamma (IFNγ) pathways as top enriched pathways (Fig. 3C-E). IFNα belong to type I IFN family which is downstream of cGAS-STING pathway (48–51). The cGAS-STING pathway has emerged as the master mediator of inflammation in the settings of stress, tissue damage, infection and cancer by sensing microbial and host derived DNAs (collectively known as danger-associated molecules-DAMPs) and is an attractive therapeutic target against cancer (40, 52, 53). Notably, decreased *STING* expression in *SMARCA4* knockout tumors is confirmed by qRT-PCR analysis (Fig. 3F). As this expression analysis was done on bulk tumors, it was unclear which cell types are responsible for these alterations.

**Figure 3:**
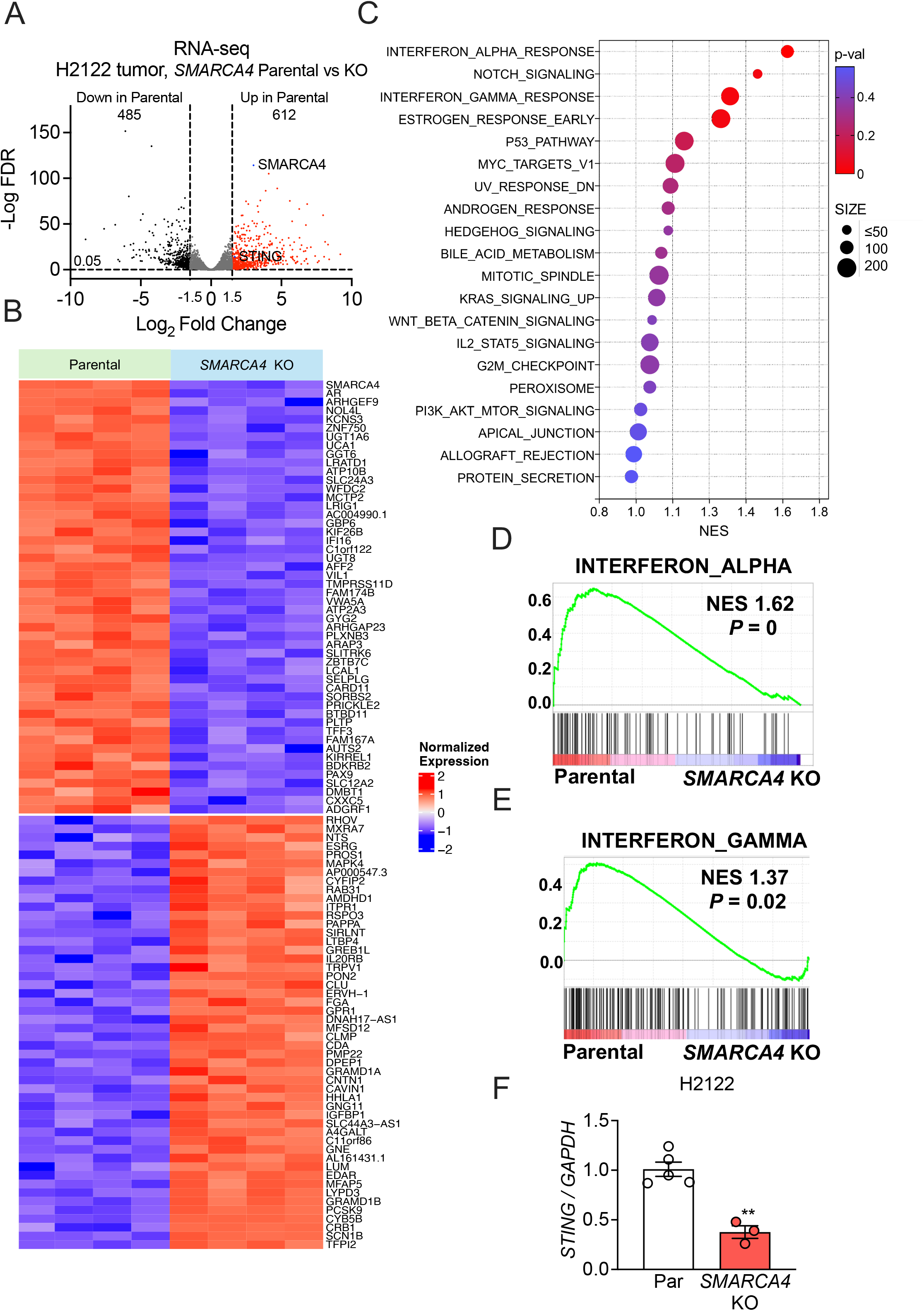
Transcriptional analysis of xenograft models identifies *SMARCA4*-dependent inflammatory response. RNA-seq analysis of parental versus *SMARCA4* KO H2122 tumors (n = 4 biological replicates). (**A**) Volcano plot displaying differentially expressed genes assessed by RNA-seq in parental and *SMARCA4* KO H2122 tumors with thresholds FDR < 0.05 and absolute log2 fold change > 1.5. (**B**) Heatmap presenting the top 50 upregulated and downregulated genes in parental tumors, with *SMARCA4* being the most significantly upregulated gene. (**C**) Gene set enrichment analysis (GSEA) of Hallmark pathways differentially enriched in parental versus *SMARCA4* KO tumors. Enrichment plots showing downregulation of (**D**) Interferon Alpha and (**E**) Interferon Gamma pathways in *SMARCA4* KO tumors. (**F**) qPCR analysis of *STING* expression in parental and *SMARCA4* KO H2122 xenograft tumors post vehicle and anti-PD1 antibody treatment, with statistical significance indicated. Data points represent mean ± S.E.M., \*\*\**p* < 0.001, \*\**p*<0.01, \**p* <0.05, two-sided unpaired t-test.

### *SMARCA4* loss results in cancer cell-intrinsic defects in expression of innate immune and inflammatory genes

SMARCA4 is a key catalytic subunit of the SWI/SNF complex which is required for proper chromatin accessibility and gene expression (54–56). Hence, we hypothesized that mutations or loss of *SMARCA4* in cancer cells could result in downregulation of *STING* and other innate immune or inflammatory genes observed above. To conclusively determine if there is a cancer cell-intrinsic dysregulation of gene expression, we performed qRT-PCR analysis of key genes such as *STING* in *in vitro* grown cancer cells. Strikingly, we noticed that *STING* expression was completely abolished in H2122 and HCC44 *SMARCA4* knockout cells (Fig. 4A-B). Next, parental and *SMARCA4* knockout H2122 and HCC44 cells were treated with STING activator cGAMP. *STING* expression is significantly increased in H2122 parental cells but not *SMARCA4* knockout cells (Fig. 4A-B). In addition to cGAMP, cGAS-STING pathway can sense cytosolic DNA to activate innate immunity (43) and initiate cytokines, such as IL-1β and IL-18 (57). In parental H2122 and HCC44 cells, the double-stranded DNA-mimetic poly(dA:dT) markedly increased expression of *STING* and *IL1β* and IL1β secretion in HCC44 (Fig. 4C-D). Consistently, IL-1β-converting enzyme, *CASPASE-1* expression is also profoundly increased by poly(dA:dT) treatment (Supplementary Fig. S6 C-D). However, *SMARCA4* knockout completely abolished the expression of these genes in response to poly(dA:dT) (Fig. 4C-D, Supplementary Fig. S6 C-D). Next, we asked if the upregulation of *STING* has actual functional consequences. One of the key downstream targets of STING pathway that plays active role in recruitment of DCs is expression of *IFNβ* (40, 53). Indeed, cGAMP treatment of parental H2122 cells potently induced expression of *IFNβ* whereas this was completely abolished in *SMARCA4* knockouts (Fig. 4B). Similarly, stimulation with poly(dA:dT) massively upregulated *IFNβ* in *SMARCA4* WT parental cells but was completely undetectable in *SMARCA4* knockouts (Fig. 4C-D). Additionally, the *IFNα* expression was highly induced by poly(dA:dT) treatment in H2122 and HCC44 parental cells but profoundly diminished in knocking out *SMARCA4* knockouts (Supplementary Fig. S6 C-D).

**Figure 4:**
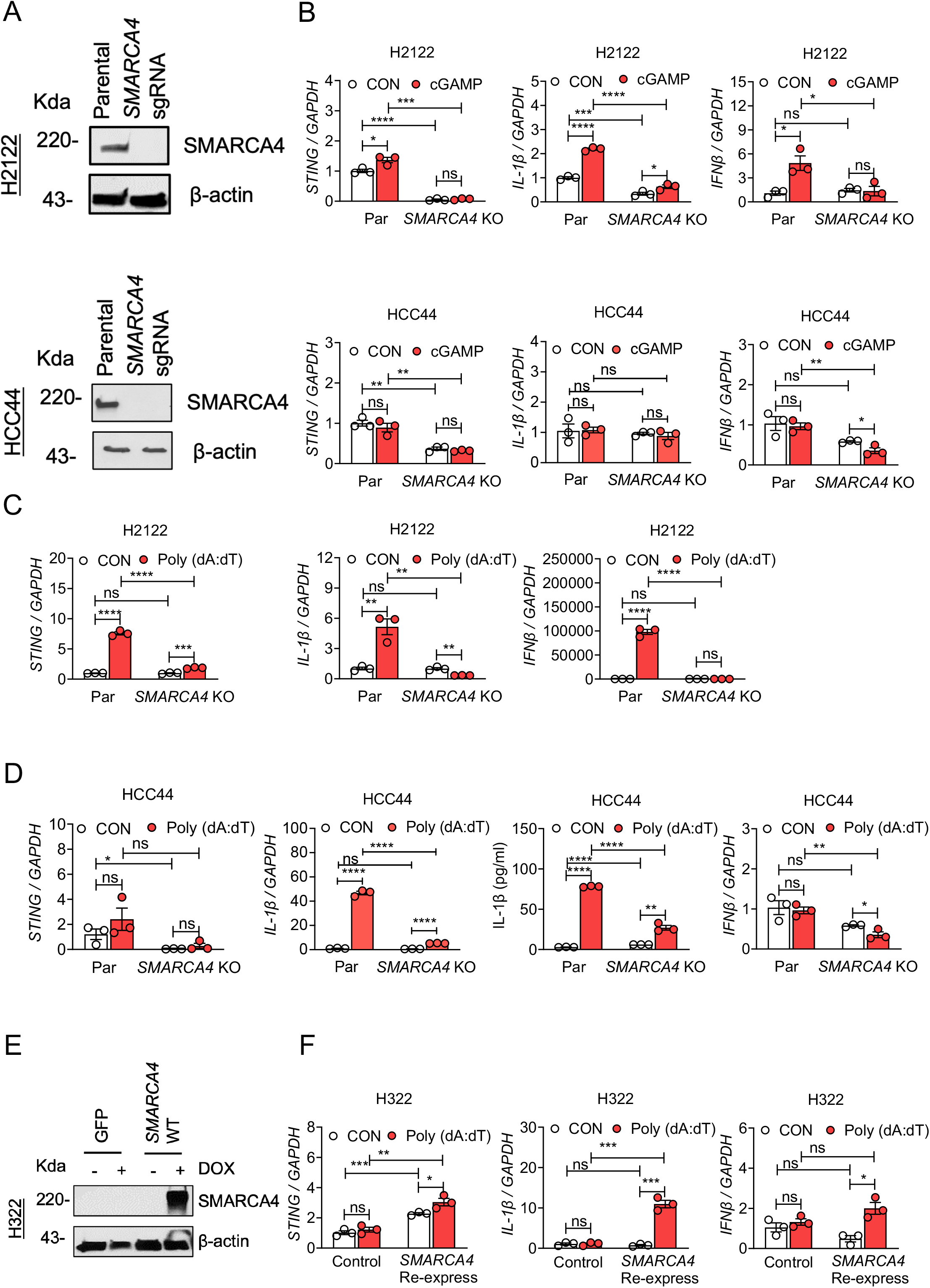
*SMARCA4* loss results in cancer cell-intrinsic defects in expression of innate immune and inflammatory genes. (**A**) Immunoblots of parental and *SMARCA4* KO H2122 and HCC44 cell lysates stained using anti-SMARCA4 and β-actin antibodies. (**B**) Expression of *STING, IL-1β* and *IFNβ* mRNA levels were evaluated in parental and *SMARCA4* KO H2122 and HCC44 cells treated with control and 10 μg/ml 2’3’-cGAMP for 6 hours (n = 3 biological replicates) by q-PCR. *STING, IL-1β* and *IFNβ* mRNA expression levels were evaluated in parental and *SMARCA4* KO (**C**) H2122 and (**D**) HCC44 cells treated with control and 5 μg/ml Poly (dA:dT) for overnight (n = 3 biological replicates) by q-PCR. HCC44 IL-1β secretion was evaluated by ELISA. (**E**) Immunoblot showing inducible expression of SMARCA4 in SMARCA4 deficient human lung cancer cell line H322 by administration of doxycycline (1 μg/ml) with GFP as control. (**F**) mRNA expression of *STING, IL-1β* and *IFNβ* by qPCR in H322 control and SMARCA4 reconstituted cell line treated with control and 5 μg/ml Poly (dA:dT) for overnight (n = 3 biological replicates). Data points represent mean ± S.E.M., \*\*\**p* < 0.001, \*\**p*<0.01, \**p* <0.05, two-sided unpaired t-test.

Type I IFNs have been widely demonstrated to be essential for cross-priming and generation of tumor specific immunity acting through DCs, potentially explaining the lack of DCs and poor immune response in *SMARCA4* mutant tumors (53, 58, 59). To definitively show the requirement of *SMARCA4*, we next sought to determine if re-expression of *SMARCA4* can rescue lack of expression of *STING* and *IL1β* in *SMARCA4* mutant cells. Hence, we inducibly re-expressed *SMARCA4* in H322, a *SMARCA4* mutant lung cancer cell line (Fig. 4E). Interestingly, re-expression of *SMARCA4* was able to potently rescue *STING* and *IL-1β* expression after poly (dA:dT) stimulation as compared to the parental cell line (Fig. 4F).

To further confirm our conclusions in an orthotopic model of lung cancer, we utilized a genetically engineered mouse (GEM) model that we have previously established termed KPS (9). Briefly, this model allows selective genetic manipulation of floxed alleles within the lung epithelium by intranasal inhalation of Adenovirus encoding Cre under the CC10 promoter resulting in inactivation of *Smarca4 and p53* and activation of mutant *Kras* (*Kras^LSLG12D/WT^, p53^fl/fl^*, *Smarca4^fl^*^/*fl*^) (Fig. 5A), following previously established protocols (60). As control, we used established KP mice (*Kras^LSLG12D/WT^, p53^fl/fl^*). Three months after Cre administration, mice were imaged with micro-CT that showed lung cancer development in the thoracic cavity of KP and KPS mice (Fig. 5B). Importantly, we were able to corroborate our previous results in this orthotopic model whereby compared to KP tumors, KPS tumors showed significantly downregulated expression of *Smarca4* and *Sting* and trends towards reduced *Ifnβ, Ifnα, Il-1β* and *Caspase-1* expression (Fig. 5C).

**Figure 5:**
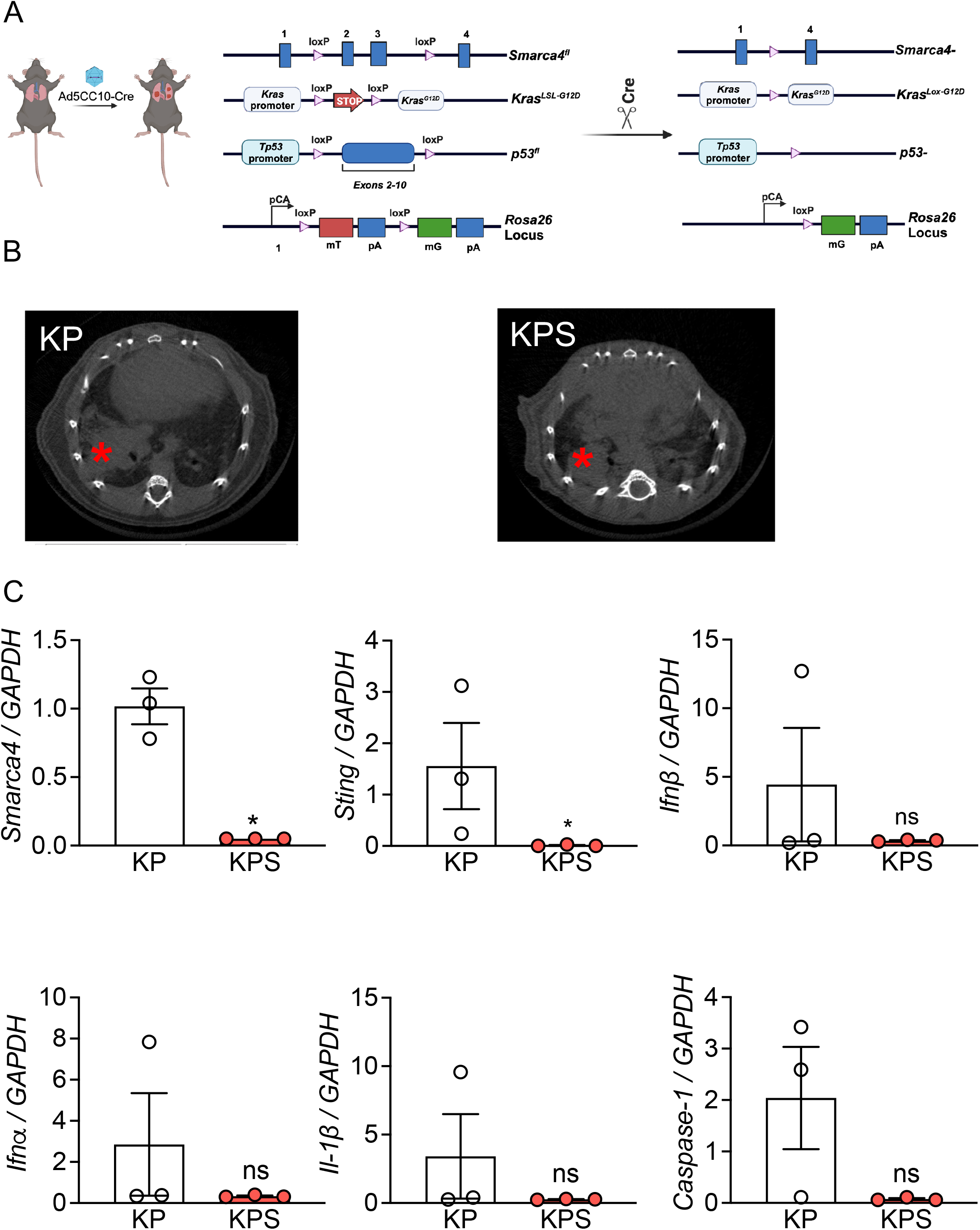
*Smarca4*-deficient GEM model tumors have diminished expression of innate immune and inflammatory genes. (**A**) Scheme of tumor induction: intranasal inhalation of adenovirus encoding CC10-Cre induces genetic recombination in airway epithelium. *Kras^G12D^* is expressed and *p53*, *Smarca4* are deleted. (**B**) microCT scanning of thoracic cavity of mice showed existence of lung cancer (marked with *). (**C**) qPCR expression of *Smarca4, Sting, Ifnβ, Ifnα, Il-1β* and *Caspase-1* of KP and KPS tumors. Data points represent mean ± S.E.M., \*\*\**p* < 0.001, \*\**p*<0.01, \**p* <0.05, two-sided unpaired t-test.

Next, we utilized a pharmacologic approach to inactivate SMARCA4 by using a potent small molecule SMARCA4 degrader based on the PROTAC principle termed ACBI1 (14). Treatment of *SMARCA4* wild type cell lines (HCC44, H1792 and H1975) by ACBI1 completely degraded SMARCA4 protein levels compared to DMSO treated cell lines. However, as expected, no obvious SMARCA4 protein levels were found in *SMARCA4* mutant cells (Fig. 6A). Gene set enrichment analysis (GSEA) revealed that top pathways downregulated in *SMARCA4* WT (HCC44, H1792 and H1975) cells upon treatment with ACBI1 were inflammatory response, TNFα signaling via NFκB and interferon alpha (IFNα) response (Fig. 6B-C). Importantly, *IL1β* was top downregulated gene by ACBI1 treatment in *SMARCA4* WT cells (Fig. 6D). The results were confirmed by qPCR in which *IL1β* expression was significantly reduced by SMARCA4 deficiency (Fig. 6E). Consistently, IL-1β-converting enzyme, *CASPASE-1* expression was also inhibited by SMARCA4 deficiency (Fig. 6F).

**Figure 6:**
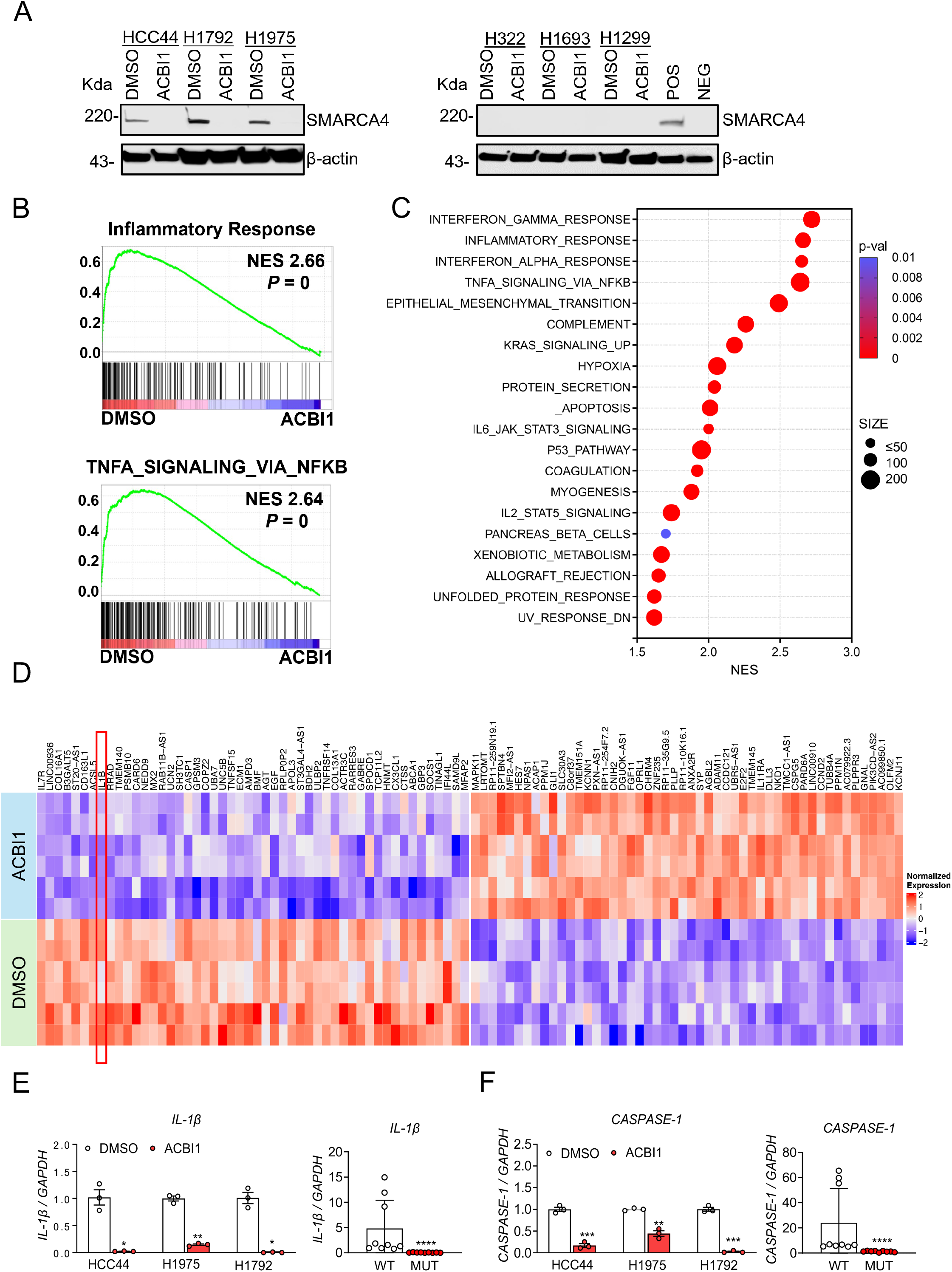
SMARCA4 degradation abrogates cancer cell-intrinsic expression of innate immune and inflammatory genes. (**A**) *SMARCA4* WT cells (HCC44, H1792 and H1975) and SMARCA4 mutant cells (H322, H1693 and H1299) were treated with DMSO control or 1 μM ACBI1 for 96 hours. Immunoblots of SMARCA4 and β-actin in *SMARCA4* WT cells (left panel) and SMARCA4 mutation cells (right panel). (**B-D**) RNA-seq analysis of *SMARCA4* WT (HCC44, H1792 and H1975) cells treated by DMSO vs 1 μM ACBI1 for 96 hours. (**B**) GSEA curves showing upregulation of the inflammatory response and TNFα_Signaling_via_NFκB pathway in DMSO treatment. (**C**) Gene set enrichment analysis (GSEA) of hallmark top enriched pathways in parental tumors. Inflammatory response and Interferon_Alpha response are at the top positions. (**D**) Heatmap presenting the top 50 upregulated and downregulated genes in DMSO treatment with *IL-1β being* the one of significantly upregulated genes. qPCR analysis of (**E**) *IL-1β* and (**F**) *CASPASE-1* expression from SMARCA4 WT cells (HCC44, H1792 and H1975) treated with DMOS vs 1 μM ACBI1 and SMARCA4 WT cells (HCC44, H1792 and H1975) vs SMARCA4 mutation cells (H322, H1693 and H1299). Data points represent mean ± S.E.M., \*\*\**p* < 0.001, \*\**p* < 0.01, \**p* < 0.05, two-sided unpaired t-test.

Taken together, these results gained through genetic and chemical biology approaches firmly establish the requirement of *SMARCA4* for basal as well as inducible expression of *STING* and *IL1β*, and broadly for expression of innate immune genes in a cancer cell-intrinsic manner.

### NF-κB transcription factor binding motif is enriched at enhancers that lose chromatin accessibility in *SMARCA4* deficient cells

Next, we asked how loss of *SMARCA4* affected expression of above-described genes. As SWI/SNF complexes actively remodel nucleosome DNA packaging, we performed Assay for Transposase-Accessible Chromatin followed by Sequencing (ATAC-Seq) upon treatment with DMSO and ACBI1 in *SMARCA4* WT cells, H1975 and HCC44. We used chemical biology approach as we can degrade SMARCA4 rapidly and assess primary molecular events that will likely help us decipher the direct actions of SMARCA4. As expected, degradation of SMARCA4 resulted in genome-wide reduction in chromatin accessibility at more than 20,000 sites in H1975 and HCC44 (Fig. 7A-B). When we examined the distribution of lost chromatin peaks, we found the majority of the peaks to be mapped to distal intergenic regions, followed by intronic regions (Fig. 7A). Many of these sites coincided with enhancers (Fig. 7A-C). These changes at distal intergenic and introns by ACBI1 are consistent with SWI/SNF complexes localizing at enhancers to regulate gene expression (54, 61, 62). Hence, we next interrogated chromatin accessibility changes at putative enhancers by performing integrative analysis and identifying genomic loci having both ATAC-seq and H3K27Acetyl ChiP-Seq peaks. We then performed pathway analysis of the genes associated with those enhancers that lost chromatin accessibility upon SMARCA4 degradation. Pathway enrichment analysis from MsigDB revealed two prominent pathways, TNF-alpha Signaling via NF-kB and Inflammatory Response were associated with lost chromatin accessibility (Fig. 7D). Enhancers are associated with specific transcription factors, hence we asked what transcription factor binding motifs were enriched in the genomic loci of enhancers in these two pathways. Interestingly, the Nuclear Factor-kappa B (NF-κB, also known as RelA or p65) motif was by far the most frequent and significantly observed sequence in the enhancers with lost accessibility (Fig. 7E). NF-κB transcription factor is a well-known and extensively studied regulator of expression of inflammatory genes (63–65). Thus, our data suggests that NF-kB and SMARCA4 could occupy the same regulatory enhancers and cooperate to drive expression of innate immune and inflammatory genes. To prove this, we next performed CUT&RUN using NF-κB and SMARCA4 antibodies and determined chromatin occupancy by these two proteins in H1975 cell line treated with DMSO or ACBI1. As expected, acute SMARCA4 degradation by ACBI1 resulted in significant reduction in chromatin accessibility and chromatin occupancy by SMARCA4 on an enhancer associated with *STING* (Fig 7F). Importantly, NF-κB is also co-localized to the same enhancer in DMSO treated cells but profoundly lost upon SMARCA4 degradation (Fig. 7F). These results strongly suggest the existence of a functional interplay between SMARCA4 and NF-κB that is critical to drive cancer cell-intrinsic expression of innate immune and inflammatory genes.

**Figure 7:**
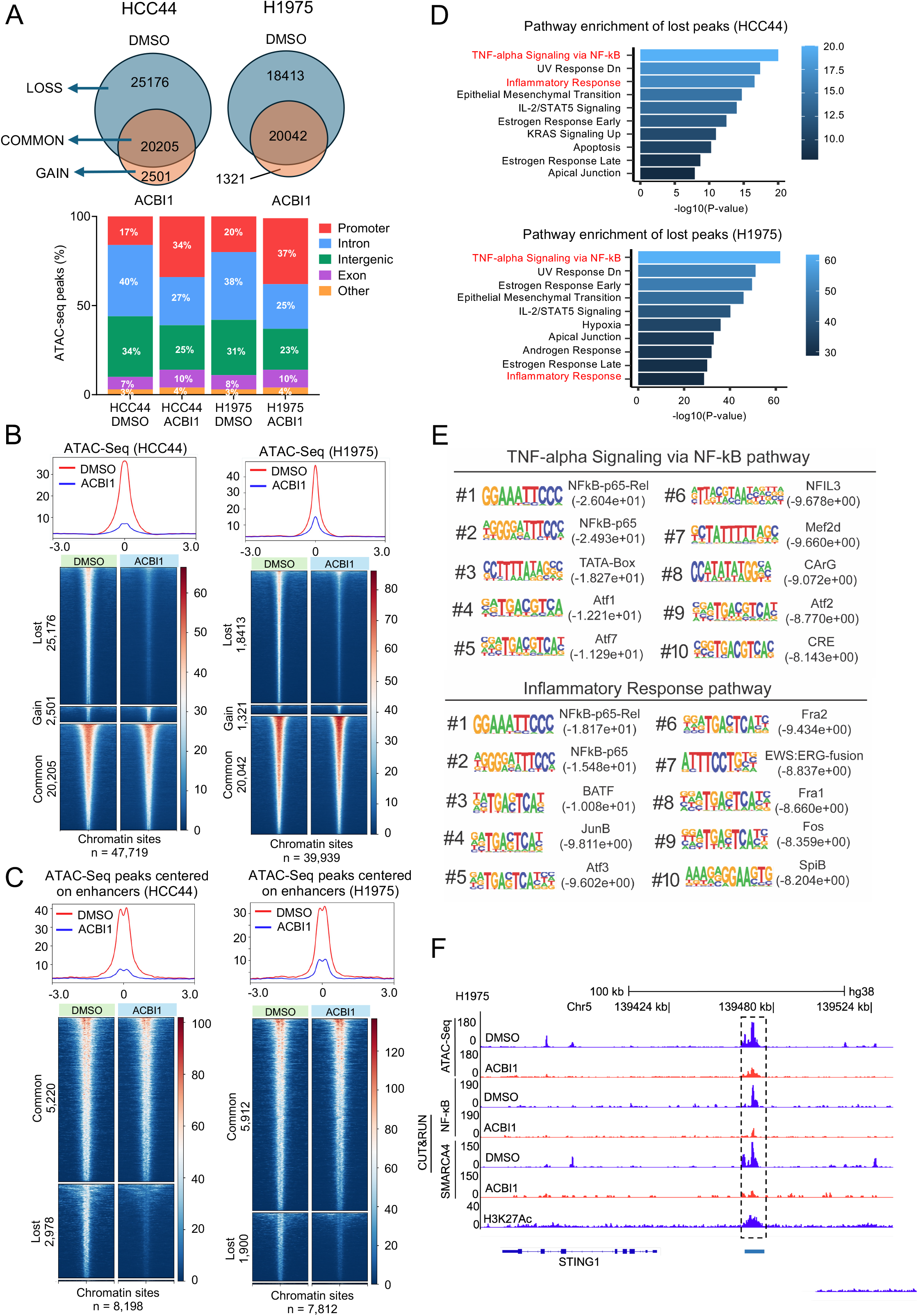
Loss of SMARCA4 abrogates chromatin accessibility at enhancers associated with innate immune, inflammation and are co-occupied by NF-κB. *SMARCA4* WT cells (HCC44 and H1792) were treated with DMSO control or 1 μM ACBI1 for 96 hours. (**A**) Venn diagrams illustrating alterations in chromatin accessibility upon ACBI1 treatment along with annotation of ATAC peaks distribution across the genomic regions. (**B**) ATAC-Seq read density heatmaps for total peaks with DMSO and ACBI1 treatment in HCC44 and H1975. (**C**) ATAC-Seq read density heatmaps for enhancer peaks with DMSO and ACBI1 treatment in HCC44 and H1975. (**D**) Pathway enrichment analysis for lost peaks in HCC44 and H1975. (**E**) Examination of transcription factor motifs on enhancers with lost accessibility as identified in (**D**). **(F)** Genome browser tracks of chromatin accessibility at *STING* loci (n = 2 biological replicates) in H1975 cells, following with representative integrative genome viewer tracks of views of the SMARCA4 and NF-κB CUT&RUN peaks on the *STING* genomic locus. H3K27Ac peaks derived from ChIP-Seq denote putative enhancers. The overlapping peaks with differential intensity in the ACBI1 treatment group are highlighted.

In conclusion, our results demonstrate that loss of *SMARCA4* in cancer cells results in profound reprograming of enhancer landscape leading to reduced expression of numerous genes including prominent regulators of innate immunity and inflammation. These transcriptional alterations are expected to drive the compromised infiltration of *SMARCA4* mutant tumors with immune cells such as DCs, culminating in an immune cold environment and lack of response to checkpoint inhibitors.

## Discussion

*SMARCA4* mutant lung cancer has emerged as a distinct clinicopathological entity with dismal prognosis and poor response to immunotherapy (6, 14, 15). In additional to primary resistance, *SMARCA4* is now implicated in acquired resistance to immunotherapy (17), suggesting a strong cancer-cell intrinsic mechanism whereby *SMARCA4* loss gives fitness advantage to tumors to escape anti-tumor immunity. In addition to lung cancer, *SMARCA4* is frequently altered across most solid tumors including endometrial carcinoma (10%), esophageal (6%), stomach (5%), bladder (5%) with a long tail of 1-2% mutation across many other solid tumors (cBioportal compilation) (66). Due to the dismal prognosis and large patient populations affected, understanding mechanisms of resistance and targeting *SMARCA4* mutant tumors is a large unmet medical need.

In this study, we provide a cancer cell-intrinsic mechanistic basis for poor response of *SMARCA4* mutant tumors to immunotherapy. Specifically, we demonstrate a mechanism by which *SMARCA4* loss leads to profound epigenetic dysregulation that primarily involves enhancers resulting in transcriptional deregulation. Part of this dysregulation involves well-studied and validated modulators of the innate immune system with roles in anti-tumor immunity such as STING and type I IFNs.

We have utilized several orthogonal and complementary model systems to show robust and reproducible molecular events associated with *SMARCA4* loss. While model-specific cellular and molecular alterations are evident, we have primarily focused our investigations on recurring themes that we observed across several model systems which are more likely to be of functional relevance. In this instance, we have observed reduced response to anti-PD1 treatment and decreased infiltration of DCs and CD4+ T cells in H2122 *SMARCA4* knockout xenograft tumors in immune-humanized as well as FM471 *Smarca4* knockout syngeneic model systems. Interestingly, DCs have emerged as a critical component of anti-tumor immunity with roles at each step of the cancer-immunity cycle including their traditionally well-known role in antigen cross-presentation at the priming phase, co-stimulatory molecule expression, secretion of cytokines that activate and stimulate migration of T cells during immunotherapy (31–36). In mouse models, DCs are necessary for therapeutic efficacy of PD1/PDL1 blockade, underscoring their essential role (36, 67). Further, activated CD4+ T cells can program DCs to boost molecular programs that are important for optimal anti-tumor immunity (68). cDC1 are primed and licensed by CD4+ T cells to induce anti-tumour immunity (69). Moreover, cDC1 are essential during the active phase of anti-PD1 treatment (70). Taken together, our observation of significant reduction in DCs, including cDC1, and CD4+ T cell is most likely responsible for the poor anti-tumor immune response of *SMARCA4* mutant tumors.

Type I IFNs and inducers of IFN response such as STING are central regulators of innate immune sensing of tumors and activators of DCs (32, 37). Using genetic loss-of-function and gain-of-function approaches as well as pharmacologic tools, we demonstrate the requirement of *SMARCA4* for proper expression of STING, type I IFNs at basal and stimulated levels. By performing epigenomic profiling, we identified deregulation of enhancers associated with innate immune genes and inflammatory genes in *SMARCA4* deficient states. Our finding of enhancer deregulation is consistent with previous observations that showed loss of SWI/SNF activity mostly alters enhancer targeting (55, 61, 71, 72). Importantly, our integrative analysis strongly suggests NF-κB transcription factor and SMARCA4 are colocalized on enhancers of genes such as *STING* in cancer cells and regulate its expression. These data are in agreement with multiple studies demonstrating that STING pathway senses cytosolic DNA and cGAMP to induce type I IFN and inflammatory cytokine genes expression via transcription factor NF-κB (73–78). Taken together, we provide a working model whereby *SMARCA4* deficiency leads to enhancer shut down at critical innate immune genes and inflammatory cytokines dampening the recruitment of DCs and T cells culminating in an immune-cold phenotype and lack of response to immunotherapy (Fig. 8). While our findings are highly interesting, a number of questions remain open for future investigations. The detailed functional interplay including direct physical interaction, dynamics of recruitment and cooperativity between SMARCA4 and NF-κB are active areas for future studies. While we have focused our investigations mainly on cancer cell-intrinsic dysregulations, it is likely that dysfunctions exist within the immune cells in *SMARCA4* mutant tumors and need to be systematically investigated. Additionally, *SMARCA4* mutant tumors have distinct metabolic features including heightened oxidative phosphorylation(9). To date it is unknown if and how these metabolic alterations affect anti-tumor immunity of *SMARCA4* mutant tumors. Lastly, a series of recent reports have identified role of SWI/SNF in various aspects of immune regulation. The Hargreaves lab showed that inhibition of SWI/SNF complex disrupts activation of gene expression in response to bacterial endotoxin in macrophages with implications in enhancer deregulation (72). The core epigenomic dysregulation observed in this report are similar to our observations suggesting conserved mechanisms. Interestingly, a recent publication demonstrated how loss of *ARID1A*, a subunit of a particular type of SWI/SNF complex termed canonical BAF complex, is associated with enhanced immunotherapy response and increased cGAS-STING pathway activation (79) which is in stark contrast to our findings of *SMARCA4* mutation. It is critical to note that there are three distinct subtypes of the SWI/SNF complex, canonical BAF (BRG-/BRM-associated factor) complex, polybromo-associated BAF (PBAF) or non-canonical BAF complexes with distinct functional roles (80). ARID1A exists only in the canonical BAF complex, whereas SMARCA4 is the catalytic subunit in all three subunits (80). Thus, the biochemical and epigenomic consequences of loss of ARID1A and SMARCA4 are expected to be divergent. Taken together, these reports from the Hargreaves, Adelman and our report suggest distinct roles for each subtype of SWI/SNF complex including potential antagonistic or competitive functions at various genomic loci. Importantly, another report has shown that ARID1A interacts with mismatch repair (MMR) protein MSH2 and its loss is correlated to microsatellite instability and increased anti-tumor immunity (81) further highlighting its distinctiveness.

**Figure 8:**
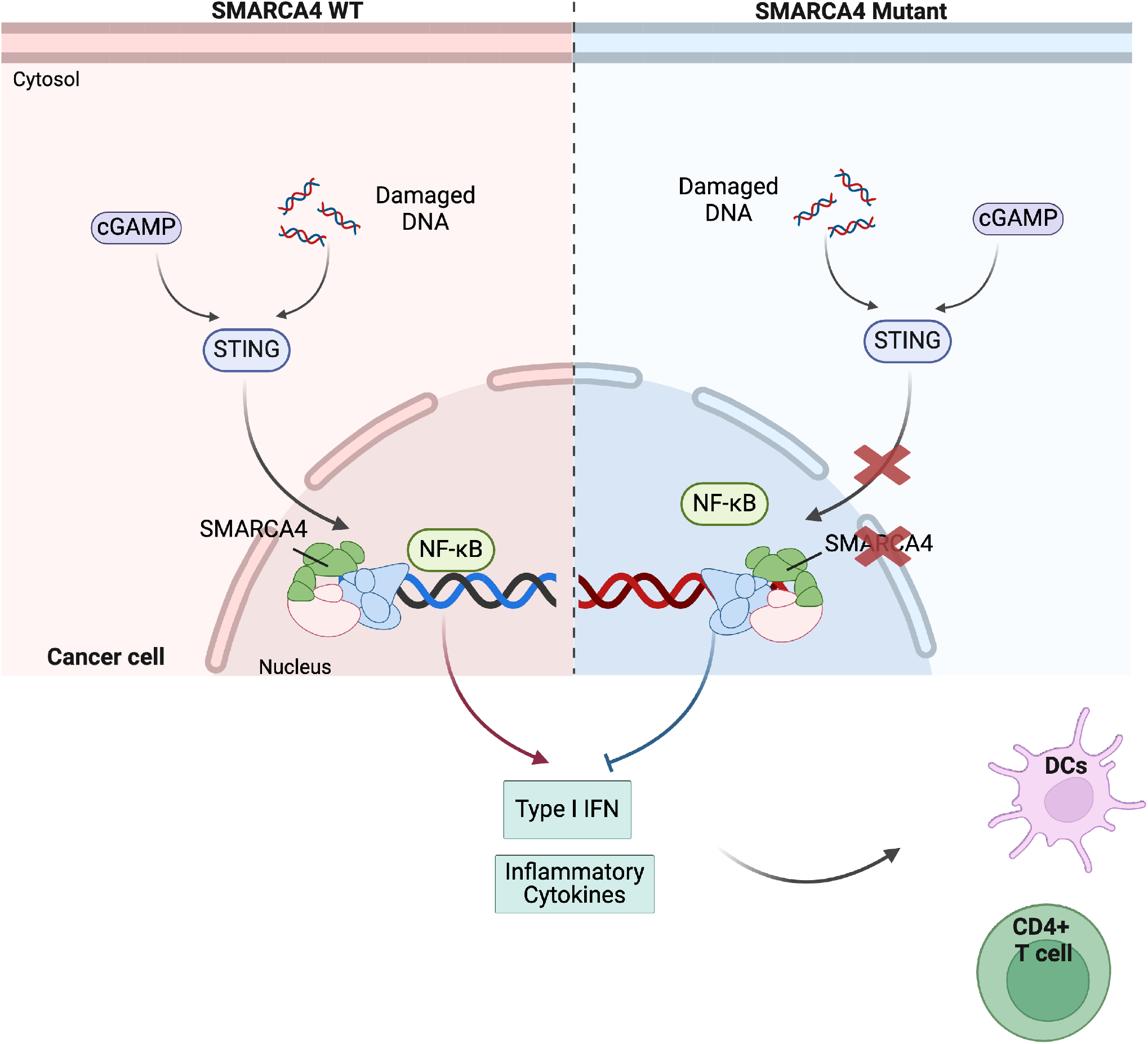
Schematic summarizing working model. In *SMARCA4* WT (left) cancer cells, cytosolic DNA sensing can be efficiently transmitted into transcriptional responses involving STING, type I IFN and inflammatory cytokines leading to recruitment of DCs and T cells. In contrast, in *SMARCA4* mutant cells (right), defective transcriptional responses prevent efficient recruitment of immune cells into the tumor microenvironment.

In conclusion, our study provided the mechanistic basis for resistance of *SMARCA4* mutant tumor to immune checkpoint blockade therapy. Based on our findings, therapeutic efforts to boost DCs or STING signaling might help overcome poor response of *SMARCA4* mutant tumors to immunotherapy.

## Author contributions

Y.W. designed experiments, interpreted the data and performed most of the experiments with assistance from I.M., M.Q., S.K. and Y.H.. Y.W. wrote the manuscript and formatted figure with assistance from M.Q.. M.Q., Y.X. and J.W. performed the computational analyses for ATAC-seq, RNA-seq and CUT&RUN. J.W. provided comments and participated in computational analysis. Y.L. conceived the work, designed the studies, supervised the project, and wrote the manuscript.

## Acknowledgements

We thank the MD Anderson Cancer Center (MDACC) Core facilities, including the Advanced Technology Genomics Core, Advanced Cytometry & Sorting Facility, DVMS Veterinary Pathology Services, and Small Animal Imaging Facility, Shan Jiang for assistance in maintenance in mouse colonies. We also thank Dr. Jack Roth, Dr. Lihui Gao and other members of the Department of Thoracic Surgery-Research section and Department of Genomic Medicine for valuable comments during the progress of this project. We are immensely grateful for Dr. Humam Kadara for providing FM471 cell line. This study was supported by The University of Texas MD Anderson start-up funds (YL). This work is also in part supported by NIH grants R01CA272945 (YL) and R37CA251629 (YL).

## Code availability

The data were analyzed by using publicly available code.

## Materials and Methods

### Ethics statement

Animal studies were carried out according to protocols approved by the University of Texas MD Anderson Cancer Center (UTMDACC) Institutional Animal Care and Use Committee. Mice were fed commercial rodent diet (PicoLab Rodent diet 5053 from Labdiet) and water *ad libitum*. All mice were kept in a specific pathogen free vivarium at the MD Anderson Cancer Center mouse facilities. Mice were kept in a 12-h light/12-h dark cycle as commonly used and housed at 18–23 °C with humidity of 50–60%.

### Humanized mice generation

Mice humanization were generated described previously (45). Female 3-4 weeks old NOD.Cg-*Prkdc^scid^ Il2rg^tm1Wjl^*/SzJ (NSG) mice source obtained from The Jackson Laboratory. Mice were housed in microisolator cages under specific pathogen-free conditions in a dedicated humanized mice room in the animal facility at The University of Texas MD Anderson Cancer Center. Mice were provided with autoclaved acidified water and fed a specialized diet known as the Uniprim diet.

Human umbilical cord blood units for research were sourced from MD Anderson Cord Blood Bank under an Institutional Review Board (IRB)-approved protocol (Lab04-0249). The cord blood bank acquires umbilical cord blood through voluntary donations from mothers following informed consent under the institutional approved IRB protocol. Cord blood bank collects human cord blood on a daily basis from several Houston-area hospitals like Memorial Hermann Hospital, St. Joseph Medical Center, and the Woman’s Hospital of Texas, etc. Fresh cord blood units were promptly transported to the research lab within 24 hours of collection for HLA typing at MD Anderson HLA typing core facility. The cord blood was diluted with phosphate-buffered saline at a 1:3 ratio, and mononuclear cells were isolated by using density-gradient centrifugation with Ficoll medium. The isolated mononuclear cells were then directly utilized for CD34+ enrichment procedures.

After mononuclear cells were separated from human umbilical cord blood, CD34+ hematopoietic stem cells were isolated using a direct CD34+ MicroBead kit (Miltenyi Biotec). 3-4 weeks old NSG mice were irradiated with 200 cGy using a ^137^Cs gamma irradiator. Over 90% pure freshly isolated CD34+ hematopoietic stem cells were injected intravenously, 24 hours after irradiation, at a density of 1-2 × 10^5^ CD34+ cells/mouse. The engraftment levels of human CD45+ cells were determined in the peripheral blood at 6 weeks post CD34+ injection by flow cytometric quantification, as well as other human immune populations. Mice with over 25% human CD45+ cells were considered humanized (Hu-NSG mice). In-depth analysis of peripheral blood for human immune cell subpopulations, including CD45+, CD3+, CD4+, and CD8+ T cells, B cells, NK cells, and lineage-negative cells, was performed using a 10-color flow cytometry panel at weeks 6 post CD34+ engraftment. Hu-NSG mice from different cord blood donors with varying levels of engraftment were randomized into every treatment group in all the experiments. All hu-NSG mice were verified for humanization before tumor implantation.

### Genetically engineered mouse model (GEMM) of lung adenocarcinoma

We have established novel inducible lung cancer mouse models, termed KPS, driven by loss of SMARCA4 in combination with *p53* loss and *Kras* mutations (*<u>K</u>ras^LSLG12D/WT^, <u>p</u>53^fl/fl^, <u>S</u>marca4^fl/f)^*(9). Briefly, *Smarca4^flox/flox^* mice with *loxp* sites flanking exons 2 and 3 of mouse *Smarca4* gene were crossed with previously described KP (*K-ras^LSL-G12D/WT^*; *p53^flox/flox^*) mice (60, 82). KP and KPS mice were backcrossed to the C57BL/6 background for 10 generations to achieve a pure genetic background. Ad5CC10-Cre virus at a viral titer of 10^12^ particles per ml (pt/ml) was purchased from The University of Iowa Viral Vector Core Facility. 5-8 weeks KP and KPS mice were intratracheally injected with 50 μl of 4 x 10^10^ pt/ml Ad5CC10-Cre virus. MicroCT scan was performed three months post-virus injection to assess lung tumor development.

### Micro-CT scans of mice

A Bruker Sky Scan 1276 (Bruker, Kontich, Belgium) was used for data-acquisition in prone position under isoflurane inhalation anaesthesia (tube voltage 70 kV, tube current 200 μA, 30 μm effective pixel size) with Retro-active respiratory gating (i.e. synchronization of acquisition of micro-CT projections with a timepoint in the respiratory cycle of the individual mouse). Scanning took approximately ∼3.5 minute. Respiratory monitoring was performed a camera and tape to monitor the respiratory sweep. Images were reconstructed and assessed at a constant window width/window level (0 - 0.025).

### Anti-PD1 treatment in Hu-NSG xenograft and syngeneic tumor models

1 x 10^7^ H2122 parental or *SMARCA4* KO cells in 200 μl PBS:Matrigel (1:1) mix were injected subcutaneously into female hu-NSG mice 6-8 weeks post CD34 engraftment. When tumor size reached approximately 200 mm^3^, 200 μg anti-human PD1 antibody (Pembrolizumab, Keytruda) or corresponding isotype control antibody were delivered intraperitoneally in 100 μl PBS to each mouse twice a week for 2-3 weeks.

For the FM471 model, 1 x 10^7^ FM471 parental or *Smarca4* KO cells in 200 μl PBS:Matrigel (1:1) mix were injected subcutaneously into female 8 weeks old C57BL/6 mice. Treatment with 200 μg rat anti mouse PD-1 IgG2a antibody (RMP1-14, Bio X Cell) or corresponding isotype control antibody in 100 μl PBS was administered to each mouse 3 time a week. The endpoint was determined when the tumor size decreased to approximately 200 mm³.

The length (L) and width (W) of the tumor were measured using calipers. The tumor size was calculated according to the following formula: L*W^2^/2.

### Cell culture

H2122, HCC44, H1792, H1975, H1693, H1299 and H322 were purchased from the American Type Culture Collection (ATCC). HCC44 was purchased from DSMZ. H322 was from Sigma. These cells were cultured in RPMI-1640 medium supplemented with 10% heat-inactivated fetal bovine serum and 1% penicillin–streptomycin at 37 °C with 5% CO_2_.

FM471 (MDA-F471) cell line (47) was kindly provided by Dr. Humam Kadara (The University of Texas MD Anderson Cancer Center, TX). These cells were derived from lung adenocarcinomas of C57/BL6 mice subjected to tobacco smoke and possess *Kras* and *p53* mutations. The cells are cultured in DMEM/F-12 medium supplemented with 10% heat-inactivated fetal bovine serum and 1% penicillin–streptomycin at 37 °C in a humidified incubator with 5% CO_2_. Routine Mycoplasma testing was performed using Mycoplasma Detection Kit.

cGAMP (10 μg/ml; InvivoGen) and poly(dA:dT) (5 μg/ml; InvivoGen) were transfected into cells using Lipofectamine 3000 (Life Technologies) diluted in Opti-MEM, according to the manufacturer’s instructions. ACBI1 (1 μM) was diluted in cell medium to treat cells. cGAMP and poly(dA:dT) were all reconstituted in ddH2O. ACBI1 was dissolved in DMSO. Equivalent amounts of ddH2O and DMSO were used for vehicle controls as appropriate.

### The generation of knockout cell lines

Human lung cancer cell lines, H2122 and HCC44, deficient in *SMARCA4* were generated by CRISPR-Cas9. sgRNAs were designed with

CRISPick (https://portals.broadinstitute.org/gppx/crispick/public).

The sgRNA was synthetized by

ThermoFisher (https://www.thermofisher.com/us/en/home/life-science/genome-editing/crispr-libraries/trueguide-grnas.html).

Transfect sgRNA to cells with TrueGuide™ Synthetic gRNA and TrueCut™ Cas9 Protein v2 using the

Neon™ Transfection System protocol (https://assets.thermofisher.com/TFS-Assets/LSG/manuals/MAN0017058_TrueGuide_Synthetic_gRNA_UG.pdf)

Mouse lung cancer cell line FM471 deficient in *Smarca4* were generated by Advanced Cell Engineering and 3D Models Core at Baylor College of Medicine using CRISPR/Cas9.

Knockout cell lines were confirmed using immunoblotting and test mycoplasma negative.

### Immunoblotting

Cells were pelleted and lysed using RIPA buffer. Cell lysates were centrifuged at 12,000 *x g* for 20 min at 4 °C and supernatants containing the soluble proteins were collected. Protein concentration was determined using Bio-Rad Protein Assay Kit II #5000002. Subsequently, 20 - 30 μg of total protein from each sample was loaded into individual lanes. Proteins were mixed with 4X LDS buffer, boiled at 95°C for 5 minutes and separated by gradient SDS-PAGE (4-15%), followed by transfer to nitrocellulose membranes. Membranes were blocked with 5% nonfat dried milk in PBS and incubated overnight at 4 °C with primary antibodies specific to the target proteins of interest. Post-incubation, membranes were washed with PBS and incubated for 1 h at room temperature with the appropriate horseradish peroxidase-conjugated (HRP) secondary, which facilitate enhanced chemiluminescence detection. Membranes were imaged using the LI-COR Odyssey software.

### RNA isolation and quantitative real-time PCR

Total RNA isolation from cells were performed with the Qiagen RNeasy kit according to the manufacturer’s instructions. For tumor RNA isolation, tumors were homogenized by zirconium oxide beads. The homogenates were purified by Qiagen RNeasy kit.

Reverse transcription from total RNA was performed from 2 μg of total RNA using the SuperScript™ IV VILO™ Master Mix (Invitrogen-Life Technologies) according to the manufacturer’s instructions. Quantitative RT-PCR was performed with SYBR Green dye using an AriaMx Real-time PCR System (Agilent). PCR reactions were performed in triplicate and the relative amount of cDNA was calculated by the comparative delta computed tomography method using *GAPDH* as an internal control.

### Preparation of single-cell suspensions for flow cytometry

Erythrocytes in the peripheral blood were lysed with ACK lysis buffer (Thermo FisheS). Single-cell suspensions were prepared from fresh primary tumors and spleen tissues using standard procedures (83, 84). The suspensions were generated by mechanical dissociation and incubation for 20min at 37 °C with dissociation solution (100 µg/mL collagenase IV and 20 µg/mL DNase I in RPMI 1640). End the digestion with 10mM EDTA at room temperature for 5-10 min. Filtered the digested tissue suspension through 100-µm nylon filter. The remaining red blood cells were lysed using 0.5 ml RBC lysis buffer by incubation 5 min at room temperature. Single cells were obtained in FACS buffer (1 x PBS, 1mM EDTA, 2% FBS).

### Flow cytometry

H2122 tumors and spleen of hu-NSG mice were staining with following antibodies: CD45.2-AF700 (clone 2D1), CD19-PE-Cy7 (clone HIB19), CD3-PerCp-Cy5.5 (clone HIT3a), CD4-Pacific Blue (clone OKT4), CD8-APC-Cy7 (clone HIT8a), CD25-APC (clone BC96), CD45RA-PE (clone HI100), CCR7-FITC (clone G043H7), CD69-PE-eflu 610 (clone FN50), CD279(PD-1)-SB702 (clone J105), CD103-SB600 (clone B-Ly7), CD56-BV510 (clone HCD56), CD19-FITC (clone HIB19), CD3-FITC (clone HIT3a), HLA-DR-PerCp-Cy5.5 (clone L243), CD33-PE (clone WM53), CD163-APC (clone GHI/61), CD11b-PE-Cy7 (clone ICRF44).

Humanized mice overall test FACS antibodies:

Anti-mouse CD45.2-FITC (clone 104), anti-human CD45.2-AF700 (clone 2D1), anti-human CD3-PerCp-Cy5.5 (clone HIT3a), anti-human CD19-PE-Cy7 (clone HIB19), anti-human CD56-PE (clone HCD56).

FM471 tumors of C57BL/6 mice were staining with following antibodies:

CD45.2-AF700 (clone 104), CD25-BV421 (clone PC61), CD4-BV510 (clone RM4.5), CD279(PD-1)-BV711 (clone 29F.1A12), CD8-FITC (clone 53-6.7), CD3-PerCP.Cy5.5 (clone 17A2), CD62L-PE (clone MEL-14), CD19-PE-Cy7 (clone 1D3), CD44-APC (clone IM7), CD69-APC-Cy7 (clone H1.2F3), CD274(PD-L1)-BV421 (clone 10F.9G2), CD11b-BV510 (clone M1/70), CD163-BV711 (clone S15049I), CD3-FITC (clone 17A2), CD19-FITC (clone 1D3), I-A/I-E(MHCII)-PerCP.Cy5.5 (clone M5/114), CD11c-PE (clone N418), CD86-PE-Cy7 (clone GL-1), CD40-APC (clone 3/23), GR-1-APC-Cy7 (clone RB6-8C5), CD69-BUV737 (clone H1.2F3), CD127-BV421 (clone IL-7Rα), KLRG1-BV510 (clone 2F1/KLRG1), CX3CR1-BV650 (clone SA011F11), CD44-BV785 (clone IM7), CD62L-PerCP.Cy5.5 (clone MEL-14), CXCR3-PE (clone CXCR3-173), CD4-PE-Cy5 (clone RM4-5), CD103-PE-Cy7 (clone 2E7), CD45RA-BV421 (clone 30-F11), CD27-BV650 (clone LG.3A10), CCR7-BV785 (clone 4B12), CD95-PE (clone SA367H8).

Dead cell staining was carried out with LIVE/DEAD™ Fixable Blue Dead Cell Stain (Invitrogen™) at 1:1,000 dilution. For flow cytometry analysis, all events were acquired on a BD LSRFortessa X-20 analysis or Attune NxT flow cytometer (Thermo Fisher) was carried out on FlowJo v10.

### Inducible ectopic expression of SMARCA4

eGFP as control or SMARCA4 were cloned into the pInducer20 doxycycline inducible lentiviral vector (Addgene 44012). Lentivirus was produced using standard virus production methods by co-transfecting target and packaging plasmids (psPAX2 – Addgene12260 and pMD2.G-Addgene 12259) into HEK293T cells. H322 cells were transduced with 0.45 μM filtered viral particles with polybrene (8 μg/ml). After 16 h of transduction, the media was changed for fresh regular growth media. 2 mg/ml G418 selected for 48h. After selection, cells were termed stably transduced. GFP or SMARCA4 expression was induced with 48h 1 μg/ml doxycycline induction.

### RNA-seq analysis

Bowtie2-build (85) was used to index the human reference genome (hg38). Raw sequencing reads were aligned to human reference genome and transcriptome gene annotation GENCODE V44 (86) using TopHat2 v2.1.1 (87). HTseq-count v0.11.0 (88) was used to count the expression level of each gene.

Differentially expressed genes (DEG) analysis was carried out with R package EdgeR v3.42.4 (89) using raw read count matrices with cutoff of FDR < 0.05 and absolute log2 fold change > 1.5. Genes were ranked by the log transformed p-values in differential expression analysis and set to negative/positive values for down/up regulation respectively, pre-ranked gene set enrichment analysis (GSEA) (90) were performed on the ranked gene list to calculate the normalized enrichment score (NES) for hallmark gene sets. For heatmap generation showing differentially expressed genes, CPM values were transformed into z-score and ComplexHeatmap, an R package was used to draw heatmap (91).

### ATAC-seq analysis

ATAC-seq fastq reads were aligned to the human genome hg38 using BWA (92). SAMtools v1.9 (93) was used to convert sam file into bam format. PCR duplicates were remove using picard-tools (v.2.27.4) MarkDuplicates function with “VALIDATION_STRINGENCY=LENIENT REMOVE_DUPLICATES=true” options.

SAMtools v1.9 was used to remove reads aligning to the mitochondrial genome and incomplete assemblies and filter mapping quality using the parameter -q 30 -F 1804.

The regions of ENCODE hg38 blacklist (https://www.encodeproject.org/annotations/ENCFF356LFX/) were filtered out.

Replicate coverage files were merged using bigWigMerge function from deepTools v3.5.4 (94) for visualization in IGV browser.

ATAC-Seq broad peaks were called using MACS2 v2.1.1 (95) software using parameters “--keep-dup all” and BAMPE option. The BigWig files were generated using deeptools for visualization in Integrative Genome Viewer (IGV) (96).

Peaks identified in replicates were combined together using bedtools (97) to generate intersect peaks. Peaks were annotated using the “annotatePeaks” function from HOMER v4.8 (98) and associated gene from each peak were identified. The lost peaks, gained peaks, and common peaks were identified using bedtools intersect function. The peak overlapping analysis, Venn-diagram visualization, and peak profile barplot were generated using in-house R script (R v4.3.0).

For data normalization and visualization, the BAM files were converted to the bigWig format using the bamCoverage with RPKM normalization. The heatmaps and average profiles were generated using the plotHeatmap and plotProfile scripts from deepTools v3.5.4 (94).

Genomic Regions Enrichment of Annotation Tool (GREAT) (99) was used to collect genes associated with lost peaks. The collected genes were used to find enriched pathways using online tool Enrichr (100). Motif analysis of genes associated with pathways was done using Homer (findMotifs.pl). Enhancer peaks were identified as the intersection of ATAC-Seq peaks and H3K27ac peaks using bedtools.

### CUT&RUN

CUT & RUN was performed using the CUTANA CUT & RUN kit v4. according to manufactures’ instruction.

Briefly, 0.5x10^6^ H1975 cells per antibody/condition were harvest and washed twice with wash buffer (20mM HEPES pH 7.5, 150mM NaCl, 0.5 mM spermidine). Subsequently, the cells were incubated with activated Concanavalin A-coated magnetic beads (Cat# 21-1401, EpiCypher) overnight at 4°C on a nutator with primary antibody at a 1:50 dilution (wash buffer + 0.05% of digitonin, 2mM EDTA). Following antibody incubation, pAG-MNase was introduced to each reaction, mixed, and allowed to incubate at room temperature for 10 minutes. 100 mM CaCl_2_ was gently added to each reaction while kept on ice, followed by a 2-hour incubation at 4°C. The reaction was quenched by adding stop buffer (340 mM NaCl, 20 mM EDTA, 4mM EGTA, 50 μg/mL RNase, 50 μg/ml glycogen). *E. coli* Spike-in DNA served as an internal control. After 10 min of incubation at 37°C, cells and beads were pelleted by centrifugation and placed on magnet to collect supernatant. Supernatant was transferred to a fresh tube and DNA was purified with CUT&RUN DNA purification kit. The released DNA was quantified using Qubit. CUT&RUN libraries were constructed using the NEBNext Ultra II DNA Library prep kit according to manufacturer’s instructions. Fragment size analysis of the libraries was conducted using TapeStation. Equimolar libraries were pooled together into one tube and sequenced on NextSeq500 platform with pair-end reads (2×75bp).

FASTQ files were quality checked using FastQC v0.11.8 (https://www.bioinformatics.babraham.ac.uk/projects/fastqc/) and paired-end reads were aligned to hg38 genome using Bowtie2 v2.4.5 (85). Picard v2.23.8 (http://broadinstitute.github.io/picard/) was used to remove PCR duplicates using the “MarkDuplicates function”. SAMtools v1.9 was employed to remove reads aligning to the mitochondrial chromosome, hg38 blacklisted regions and incomplete assemblies and filter mapping quality using the parameter “-q 30 -F 1804”. The read coverage, bigwig files were created with deeptools bamCoverage (94). Replicate coverage files were merged using “bigWigMerge” function from deepTools v3.5.4 (94) for visualization in IGV browser. MACS2 v2.1.1 (95) was used to call narrow peaks using parameters “--keep-dup auto” and BAMPE option.

Peaks identified in replicates were combined together using bedtools (97) to generate intersect peaks. The lost peaks were identified using bedtools intersect function. GREAT tool (99) was used to annotate the peaks in the long-distance region.

### Data availability

The RNA-seq and ATAC-seq data have been submitted to the Gene Expression Omnibus under accession number GSE269551, GSE269618, GSE269930 and GSE269686 respectively.

GraphPad Prism 8 software was used to generate graphs and statistical analyses. Statistical significance was determined by unpaired Student’s t-test. Methods for statistical tests, the exact value of n, and definition of error bars were indicated in figure legends.

