## Supplemental materials for "*SMARCA4* mutation induces tumor cell-intrinsic defects in enhancer landscape and resistance to immunotherapy"

*Corresponding author

**Supplementary Figure 1: Validation of humanization in Hu-NSG mouse model.**

Flow cytometry results obtained from mouse peripheral blood. (**A**) Human CD45+ lymphocytes were gated out from mouse cells followed by stepwise gating for human (**B**) CD3+ T cells, CD19+ B cells, and (**C**) CD56+ NK cells subpopulations. (**D**) Analysis of humanization status in peripheral blood at 6 weeks after CD34+ HSCs implantation. (**E**) Human CD4+ and CD8+ T cells gated out from mouse spleen at the endpoint of immunotherapy experiments.

**Supplementary Figure 2: *SMARCA4* deletion and immune checkpoint blockade therapy modify tumor-infiltrating lymphocyte characterization in Hu-NSG xenograft tumors.**

Flow cytometry results obtained from dissociated subcutaneously transplanted parental and *SMARCA4* KO H2122 tumors treated with vehicle or anti-PD1 antibodies. (**A**) Percentage of CD3+ T cells out of CD45+ cells as well as CD3+ T cell subtypes. (**B**) Percentage of CD4+ T cell subtypes. (**C**) Percentage of CD8+ T cell subtypes. (**D**) Percentage of CD19+ B cells and Human Regular B cells. (**E**) NK cells and PD-1 NK cells percentage. (**F**) Percentage of MDSC, CD11b+, monocytes, M1 macrophage, M2 macrophage, M1 M2 macrophage. Data points represent mean ± S.E.M., ** $p<0.01$, * $p<0.05$, two-sided unpaired t-test.

**Supplementary Figure 3: *Smarca4* affects anti-PD1 treatment efficacy and tumor-infiltrating lymphocyte characterization in mouse syngeneic models.**

Dissociated subcutaneously transplanted parental and *Smarca4* KO FM471 tumors treated with vehicle or anti-PD1 antibodies. (**A**) Five representative tumor images for each group. (**B**) Flow cytometry plot of cDC1 gating from CD11b+CD11c+ cells. (**C**) Percentage of CD3+ T cells out of CD45+ cells. (**D**) Percentage of CD4+CD8+, CD4-CD8- T cells out of CD3+ T cells. (**E**) CD19+ B cell percentage. (**F**) Percentage of MDSC.

(**G-H**) CD11b+ cell subtype percentage. (**I**) The percentage of macrophage. Data points represent mean ± S.E.M., ***p*<0.01, **p* <0.05, two-sided unpaired t-test.

**Supplementary Figure 4: Effect of *Smarca4* on CD4+ T cell subtypes in syngeneic mouse models.**

Flow cytometry results obtained from dissociated subcutaneously transplanted parental and *Smarca4* KO FM471 tumors treated with vehicle or anti-PD1 antibodies. (**A**) Percentage of CD4+CD25+, CD4+PD1+ and CD4+CD69+ T cells. (**B**) Percentage of CD4+ T naïve (CD44-CD62L+), TEM (CD44+CD62L-) and TCM (CD44+CD62L+) cells. (**C**) Percentage of CD4+ TEMRA (CD45RA+CCR7-), TEM (CD45RA-CCR7-) and TCM (CD45RA-CCR7+) cells. (**D**) Percentage of CD4+ TRM (CD103+CD69+) and TE (KLRG1+CX3CR1+) cells. (**E**) Percentage of CD4+ TMEM (KLRG1-CXCR3+), TINT (KLRG1+CXCR3+) and TEFF (KLRG1+CXCR3-) cells. (**F**) Percentage of CD4+ MP (KLRG1-CD27+) and TE (KLRG1+CD27-) cells. Data points represent mean ± S.E.M., ***p*<0.01, **p* <0.05, two-sided unpaired t-test.

**Supplementary Figure 5: Effect of *Smarca4* on CD8+ T cell subtypes in syngeneic mouse models.**

Flow cytometry results obtained from dissociated subcutaneously transplanted parental and *Smarca4* KO FM471 tumors treated with vehicle or anti-PD1 antibodies. (**A**) Percentage of CD8+PD1+ and CD8+CD69+ T cells. (**B**) Percentage of CD8+ T naïve (CD44-CD62L+), TEM (CD44+CD62L-) and TCM (CD44+CD62L+) cells. (**C**) Percentage of CD8+ TEMRA (CD45RA+CCR7-), TEM (CD45RA-CCR7-) and TCM (CD45RA-CCR7+) cells. (**D**) Percentage of CD8+ TRM (CD103+CD69+) and TE (KLRG1+CX3CR1+) cells. (**E**) Percentage of CD8+ TMEM (KLRG1-CXCR3+), TINT (KLRG1+CXCR3+) and TEFF (KLRG1+CXCR3-) cells. (**F**) Percentage of CD8+ MP (KLRG1-CD27+) and TE (KLRG1+CD27-) cells. Data points represent mean ± S.E.M., ***p*<0.01, **p* <0.05, two-sided unpaired t-test.

**Supplementary Figure 6: STING pathway activation depends on *SMARCA4*.**

(**A-B**) Parental and *SMARCA4* KO H2122 and HCC44 cells treated with control and 10 µg/ml 2'3'-cGAMP for 6 hours (n = 3 biological replicates). *CASPASE-1* and *IFNα* mRNA expression levels were evaluated in (**A**) H2122 and (**B**) HCC44 cells by qPCR. (**C-E**) Parental and *SMARCA4* KO H2122 and HCC44 cells as well as control and SMARCA4 reconstituted H322 cells are treated with control and 5 µg/ml Poly (dA:dT) for overnight (n = 3 biological replicates). *CASPASE-1* and *IFNα* mRNA expression levels were evaluated in (**C**) H2122 and (**D**) HCC44 (**E**) H322 cells by qPCR. Data points represent mean ± S.E.M., ***$p < 0.001$, **$p<0.01$, *$p <0.05$, two-sided unpaired t-test.

**Supplementary table. Key resource table**

| Reagent or Resource | Source | Identifier |
|---|---|---|
| **Antibodies** | | |
| Pembrolizumab (anti-PD-1)-human | MDACC | KEYTRUDA 88078K |
| Anti-human-CD45.2-AF700 | Biolegend | 368514 |
| Anti-human-CD19-PE-Cy7 | Biolegend | 302216 |
| Anti-human-CD3-PerCp-Cy5.5 | Biolegend | 300328 |
| Anti-human-CD4-Pacific Blue | Biolegend | 317429 |
| Anti-human-CD8-APC-Cy7 | Biolegend | 300926 |
| Anti-human-CD25-APC | Biolegend | 302610 |
| Anti-human-CD45RA-PE | Biolegend | 304108 |
| Anti-human-CCR7(CD197) -FITC | Biolegend | 353216 |
| Anti-human-CD69-PE-eflu 610 | Thermo Fisher | 61-0699-42 |
| Anti-human-CD279(PD-1)-SB702 | Thermo Fisher | 67-2799-42 |
| Anti-human-CD103(Integrin alpha E)-SB600 | Thermo Fisher | 63-1038-42 |
| Anti-human-CD56-BV510 | Biolegend | 318340 |
| Anti-human-CD19-FITC | Biolegend | 302206 |

| *Continued* | | |
| --- | --- | --- |
| **Reagent or Resource** | **Source** | **Identifier** |
| Anti-human-CD3-FITC | Biolegend | 300306 |
| Anti-human-HLADR-PerCp-Cy5.5 | Biolegend | 307630 |
| Anti-human-CD33-PE | Biolegend | 303404 |
| Anti-human-CD163-APC | Thermo Fisher | 17-1639-42 |
| Anti-human-CD11b-PE-Cy7 | Biolegend | 301322 |
| Anti-human-CD56-PE | Biolegend | 318306 |
| Anti-mouse-CD45.2-FITC | | 109806 |
| Rat anti-mouse IgG2a isotype control | Bio X Cell | BP0089 |
| Rat anti-mouse IgG2a isotype control PD-1 (CD279) | Bio X Cell | BP0146 |
| Anti-mouse-CD45.2-AF700 | BioLegend | 109822 |
| Anti-mouse-CD25-BV421 | BD | 562606 |
| Anti-mouse-CD4-BV510 | BD | 563106 |
| Anti-mouse-CD279(PD-1)-BV711 | BioLegend | 135231 |
| Anti-mouse-CD8-FITC | BioLegend | 100706 |
| Anti-mouse-CD3-PerCP.Cy5.5 | BD | 560527 |
| Anti-mouse-CD62L-PE | BioLegend | 104408 |

*Continued*

| Reagent or Resource | Source | Identifier |
|---|---|---|
| Anti-mouse-CD19-PE-Cy7 | BD | 552854 |
| Anti-mouse-CD44-APC | BD | 559250 |
| Anti-mouse-CD69-APC-Cy7 | BioLegend | 104526 |
| Anti-mouse-CD274(PD-L1)-BV421 | BioLegend | 124315 |
| Anti-mouse-CD11b-BV510 | BioLegend | 101263 |
| Anti-mouse-CD163-BV711 | BioLegend | 155325 |
| Anti-mouse-CD3-FITC | BD | 555274 |
| Anti-mouse-CD19-FITC | BD | 553785 |
| Anti-mouse-I-A/I-E(MHCII)-PerCP.Cy5.5 | BD | 562363 |
| Anti-mouse-CD11c-PE | BioLegend | 117308 |
| Anti-mouse-CD86-PE-Cy7 | BioLegend | 105014 |
| Anti-mouse-CD40-APC | BioLegend | 124612 |
| Anti-mouse-Gr-1(Ly-6G/Ly-6C)-APC-Cy7 | BioLegend | 108424 |
| Anti-mouse-CD69-BUV737 | Thermo Scientific | 367-0691-80 |
| Anti-mouse-CD127-BV421 | BioLegend | 135023 |
| Anti-mouse-KLRG1-BV510 | BioLegend | 138421 |

***Continued***

| Reagent or Resource | Source | Identifier |
|---|---|---|
| Anti-mouse-CX3CR1-BV650 | BioLegend | 149033 |
| Anti-mouse-CD44-BV785 | BioLegend | 103059 |
| Anti-mouse-CD62L-PerCP.Cy5.5 | BioLegend | 104432 |
| Anti-mouse-CXCR3(CD183)-PE | eBioscience™ | 12-1831-82 |
| Anti-mouse-CD4-PE-Cy5 | BioLegend | 100514 |
| Anti-mouse-CD103-PE-Cy7 | BioLegend | 121426 |
| Anti-mouse-CD45RA-BV421 | Thermo Scientific | 13-0451-82 |
| Anti-mouse-CD27-BV650 | BioLegend | 124233 |
| Anti-mouse-CCR7-BV785 | BioLegend | 120127 |
| Anti-mouse-CD95-PE | BioLegend | 152608 |
| SMARCA4 (Brg1) Rabbit mAb | Cell Signaling Technology | 49360 |
| β-actin | Sigma | A2228 |
| Anti-mouse IgG, HRP-linked Antibody | Cell Signaling Technology | 7076 |
| Anti-rabbit IgG, HRP-linked Antibody | Cell Signaling Technology | 7074 |

*Continued*

| Reagent or Resource | Source | Identifier |
|---|---|---|
| NF-κB p65 (D14E12) XP® Rabbit mAb | Cell Signaling Technology | 8242 |
| **Virus Strains and Plasmid** | | |
| Ad5CC10-Cre | UI Viral Vector Core | VVC-Berns-1166 |
| psPAX2 | Addgene | 12260 |
| pMD2.G | Addgene | 12259 |
| pInducer20 doxycycline inducible lentiviral vector | Addgene | 44012 |
| BRG1/SMARCA4 CUTANA™ CUT&RUN Antibody | Epicypher | 13-2002 |
| **Chemicals and Kits** | | |
| CD34+ MicroBead kit | Miltenyi Biotec | 130-046-702 |
| DPBS 1X | Sigma | D8537 |
| Cultrex Basement Membrane Extract, PathClear (Matrigel) | R&D Systems | 3432-001 |
| RPMI 1640, Glutamine | Fisher Scientific | SH30027FS |
| DMEM/F-12 HEPES | Fisher Scientific | SH30023FS |
| DMEM, high glucose, GlutaMax, HEPES | Life Technologies | 10564029 |
| Opti-MEM I Reduced Serum Medium | Life Technologies | 31985088 |
| Tet System Approved FBS | Takara | 631368 |

| *Continued* | | |
|---|---|---|
| **Reagent or Resource** | **Source** | **Identifier** |
| Fetal Bovine Serum, qualified, heat inactivated, United States | Life Technologies | 16140071 |
| PEN-STREP 100X Solution. 100ML | Cytiva | SV30010 |
| EDTA (0.5 M), pH 8.0, RNase-free | Invitrogen | AM9261 |
| Ficoll-Paque™ PLUS Media | Cytiva | 45-001 |
| ACK lysis buffer | Thermo Fisher | A1049201 |
| 2’3’-cGAMP | InvivoGen | tlrl-nacga23 |
| Poly(dA:dT) | InvivoGen | tlrl-patn-1 |
| Lipofectamine 3000 Transfection Reagent | Invitrogen | L3000008 |
| ACBI1 | Selleck Chemicals | S9612 |
| TrueCut™ Cas9 Protein v2 | Invitrogen | A36498 |
| 1xTE Buffer | Thermo Fisher | J75793.AE |
| Neon™ Transfection System 10 µL Kit | Invitrogen | MPK1025 |
| Human IL-1 beta/IL-1F2 ELISA Kit | R&D Systems | DLB50 |
| RIPA Lysis Buffer | Boston BioProducts | BP-115 |
| NuPAGE LDS Sample Buffer (4X) | Life Technologies | NP0007 |

*Continued*

| Reagent or Resource | Source | Identifier |
|---|---|---|
| Bio-Rad Protein Assay Kit II | Bio-Rad | 5000002 |
| ChIC / CUT&RUN Kit | Epicypher | 14-1048 |
| NEBNext® Ultra™ II DNA Library Prep Kit | New England Biolabs | E7645S |
| D1000 Reagents | Agilent Technologies | 5067-5583 |
| Tape station tapes: D1000 ScreenTape | Agilent Technologies | 5067-5582 |
| NP-40 | Sigma/Roche | 11332473001 |
| Illumina Tagment DNA Enzyme and Buffer Small Kit | Illumina | 20034197 |
| MinElute Reaction Cleanup Kit | Qiagen | 28204 |
| NEBNext® High-Fidelity 2X PCR Master Mix | New England Biolabs | M0541S |
| Agencourt AMPure XP magnetic beads | Beckman Coulter | A63880 |
| Qubit dsDNA HS Assay Kit | Thermo Fisher | Q32851 |
| 4–15% Mini-PROTEAN® TGX™ Precast Protein Gels | Bio-Rad | 4561085 |
| Trans-Blot Turbo Mini 0.2 µm Nitrocellulose Transfer Packs | Bio-Rad | 1704158 |

| *Continued* | | |
|---|---|---|
| **Reagent or Resource** | **Source** | **Identifier** |
| SuperSignal™ West Pico PLUS Chemiluminescent Substrate | Thermo Fisher | 34580 |
| RNeasy Mini Kit | Qiagen | 74104 |
| SuperScript™ IV VILO™ Master Mix | Invitrogen | 11766050 |
| PowerUp™ SYBR™ Green Master Mix | Applied Biosystems™ | A25742 |
| Collagenase Type IV | Sigma-Aldrich | C4-22-1G |
| DNase I | Sigma-Aldrich | 10104159001 |
| eBioscience™ 1X RBC Lysis Buffer | Invitrogen | 00-4333-57 |
| LIVE/DEAD™ Fixable Blue Dead Cell Stain Kit | Invitrogen | Invitrogen |
| pInducer20 doxycycline inducible lentiviral vector | Addgene | 44012 |
| psPAX2 | Addgene | 12260 |
| pMD2.G | Addgene | 12259 |
| polybrene | Fisher Scientific | TR-1003-G |
| G418 (Geneticin) | Invivogen | ant-gn-1 |
| Doxycycline | Fisher Scientific | BP26535 |
| **Deposited Data** | | |
| RNA-seq of tumors | This paper | GEO: GSE269551 |

| *Continued* | | |
|---|---|---|
| **Reagent or Resource** | **Source** | **Identifier** |
| RNA-seq of cells | This paper | GEO: GSE269618; GSE269930 |
| ATAC-seq of cells | This paper | GEO: GSE269686 |
| **Mouse Models** | | |
| C57BL/6J CD45.2 | The Jackson Laboratory | 000664 |
| NOD.Cg-Prkdcscid Il2rgtm1Wjl/SzJ (NSG) | The Jackson Laboratory | 005557 |
| **qPCR Primers** | | |
| Anti-human: | | |
| *IL-1$\beta$* For | CCACAGACCTTCCAGGAGAATG | N/A |
| *IL-1$\beta$* Rev | GTGCAGTTCAGTGATCGTACAGG | N/A |
| *CASPASE-1* For | GCTGAGGTTGACATCACAGGCA | N/A |
| *CASPASE-1* Rev | TGCTGTCAGAGGTCTTGTGCTC | N/A |
| *STING (TMEM173)* For | CCTGAGTCTCAGAACAACTGCC | N/A |
| *STING (TMEM173)* Rev | GGTCTTCAAGCTGCCCACAGTA | N/A |
| *IFN$\alpha$* For | AGAAGGCTCCAGCCATCTCTGT | N/A |
| *IFN$\alpha$* Rev | TGCTGGTAGAGTTCGGTGCAGA | N/A |
| *IFN$\beta$* For | CTTGGATTCCTACAAAGAAGCAGC | N/A |
| *IFN$\beta$* Rev | TCCTCCTTCTGGAACTGCTGCA | N/A |
| *GAPDH* For | AAGGTGAAGGTCGGAGTCAAC | N/A |
| *GAPDH* Rev | ACCAGAGTTAAAAGCAGCCCT | N/A |
| Anti-mouse: | | |
| *Il-1$\beta$* For | TGGACCTTCCAGGATGAGGACA | N/A |
| *Il-1$\beta$* Rev | GTTCATCTCGGAGCCTGTAGTG | N/A |
| *Caspase-1* For | GGCACATTTCCAGGACTGACTG | N/A |

| *Continued* | | |
|---|---|---|
| **Reagent or Resource** | **Source** | **Identifier** |
| *Caspase-1* Rev | GCAAGACGTGTACGAGTGGTTG | N/A |
| *Sting (Tmem173)* For | ATGTCCAGTCCAGTTGGATGTT | N/A |
| *Sting (Tmem173)* Rev | ACAGACTGCAGAGACTTCCG | N/A |
| *Ifn*$\alpha$ For | GGATGTGACCTTCCTCAGACTC | N/A |
| *Ifn*$\alpha$ Rev | ACCTTCTCCTGCGGGAATCCAA | N/A |
| *Ifn*$\beta$ For | GCCTTTGCCATCCAAGAGATGC | N/A |
| *Ifn*$\beta$ Rev | ACACTGTCTGCTGGTGGAGTTC | N/A |
| *Gapdh* For | CATCACTGCCACCCAGAAGACTG | N/A |
| *Gapdh* Rev | ATGCCAGTGAGCTTCCCGTTCAG | N/A |
| **sgRNA Sequences** | | |
| H-SMARCA4 | GCAGCAGACAGACGAGTACG | N/A |
| M-SMARCA4-F | TCATCCCTAACCAGTGGCTG | N/A |
| M-SMARCA4-R | TCTCCTGAAGGGTGGAAGTG | N/A |
| **Software and Machine** | | |
| Bruker Sky Scan 1276 | Bruker, Kontich, Belgium | N/A |
| gentleMACS™ Dissociator | Miltenyi Biotec | 130-093-235 |
| Attune Flow Cytometers | Thermo Fisher | N/A |
| Image J | Fiji | N/A |
| Neon Transfection System | Thermo Fisher | N/A |
| Odyssey Fc Imager | LI-COR Biosciences | N/A |
| AriaMx Real-time PCR System | Agilent | N/A |
| BD LSRFortessa X-20 | BD | N/A |
| FlowJo™ v10.10 | BD | N/A |
| Qubit fluorometer | ThermoFisher | N/A |

***Continued***

| **Reagent or Resource** | **Source** | **Identifier** |
|---|---|---|
| Agilent Bioanalyzer | Agilent | N/A |
| GraphPad Prism v.10.3 | GraphPad Software | N/A |

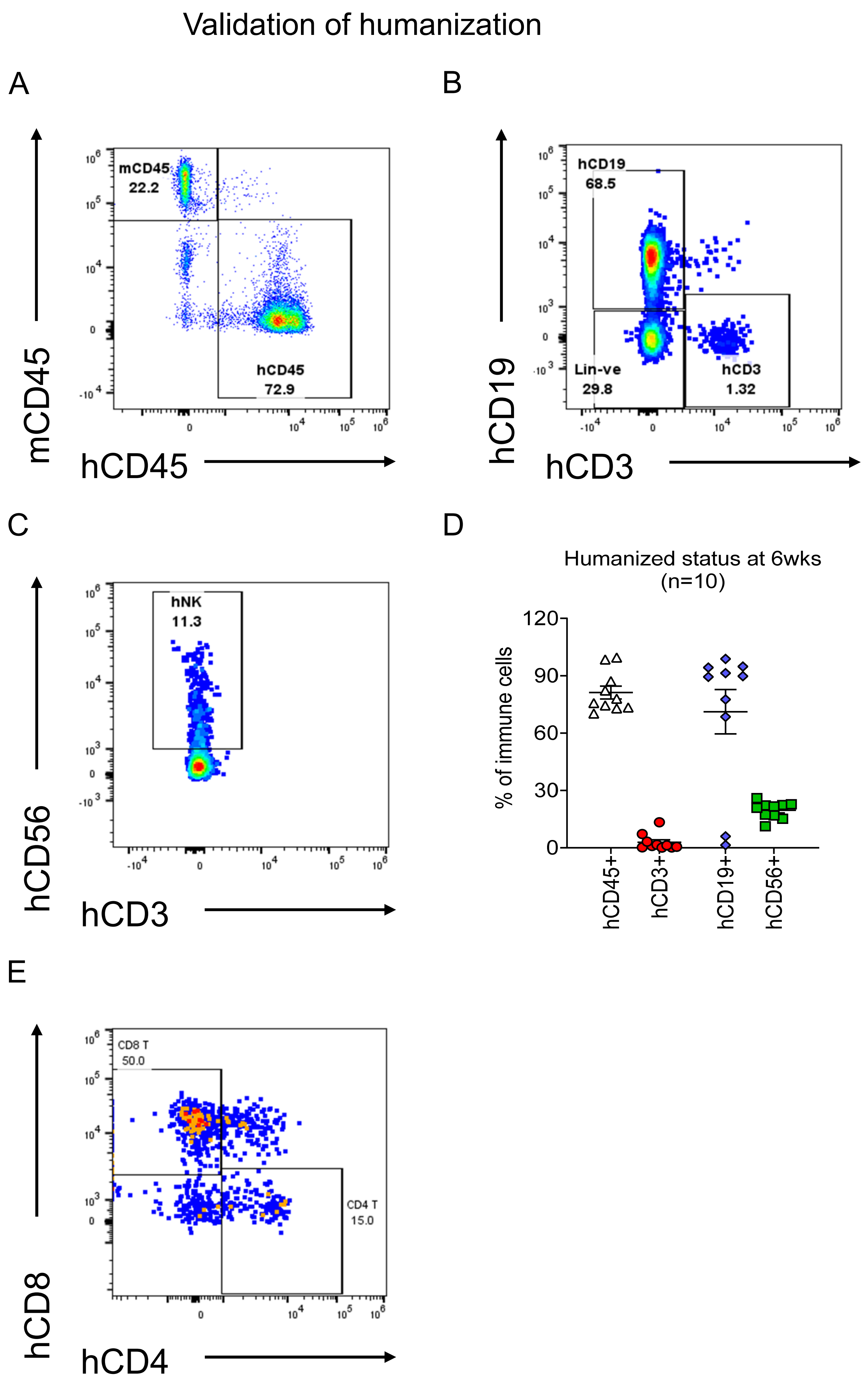


**Supplementary figure 1**

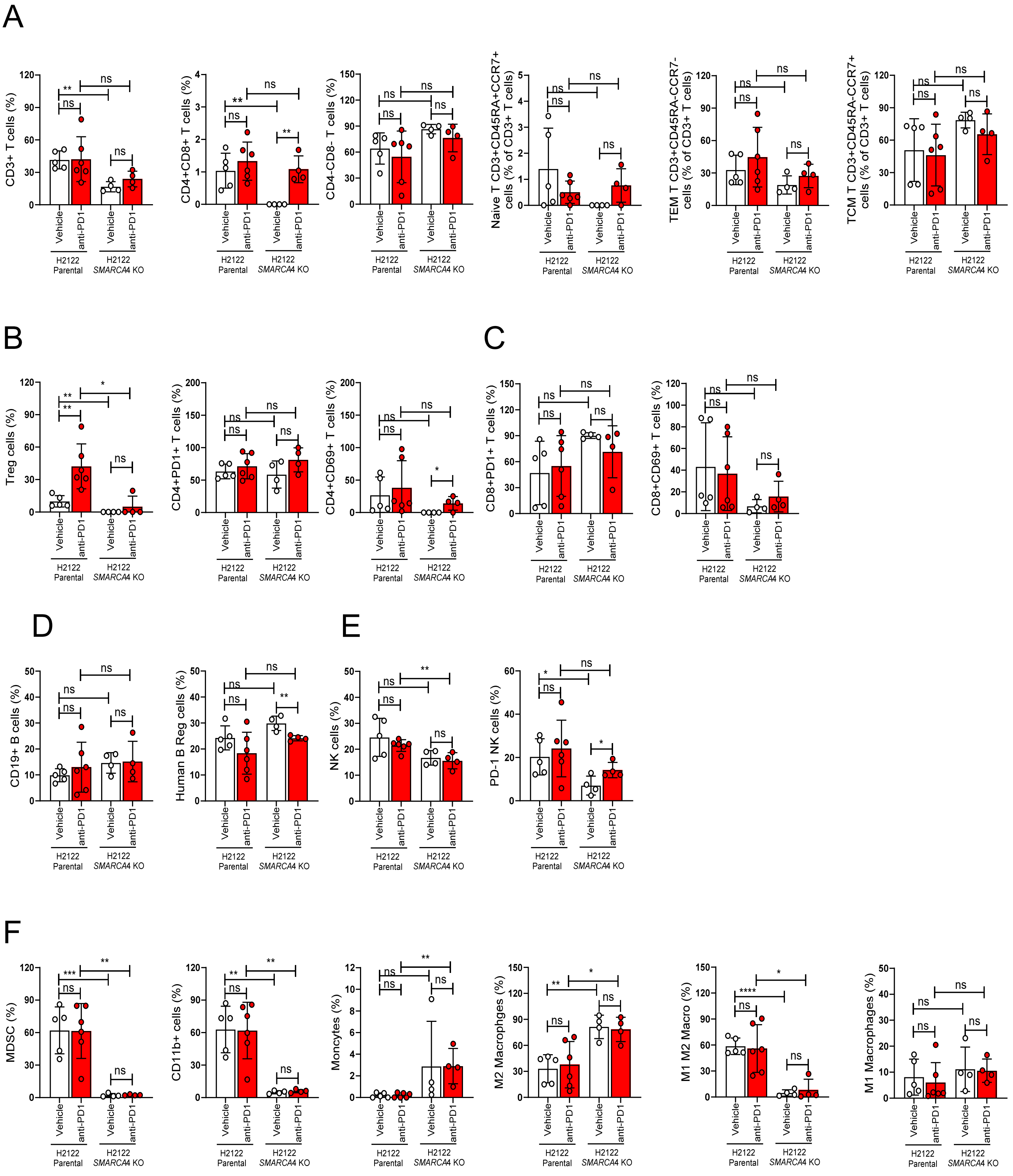


Supplementary figure 2

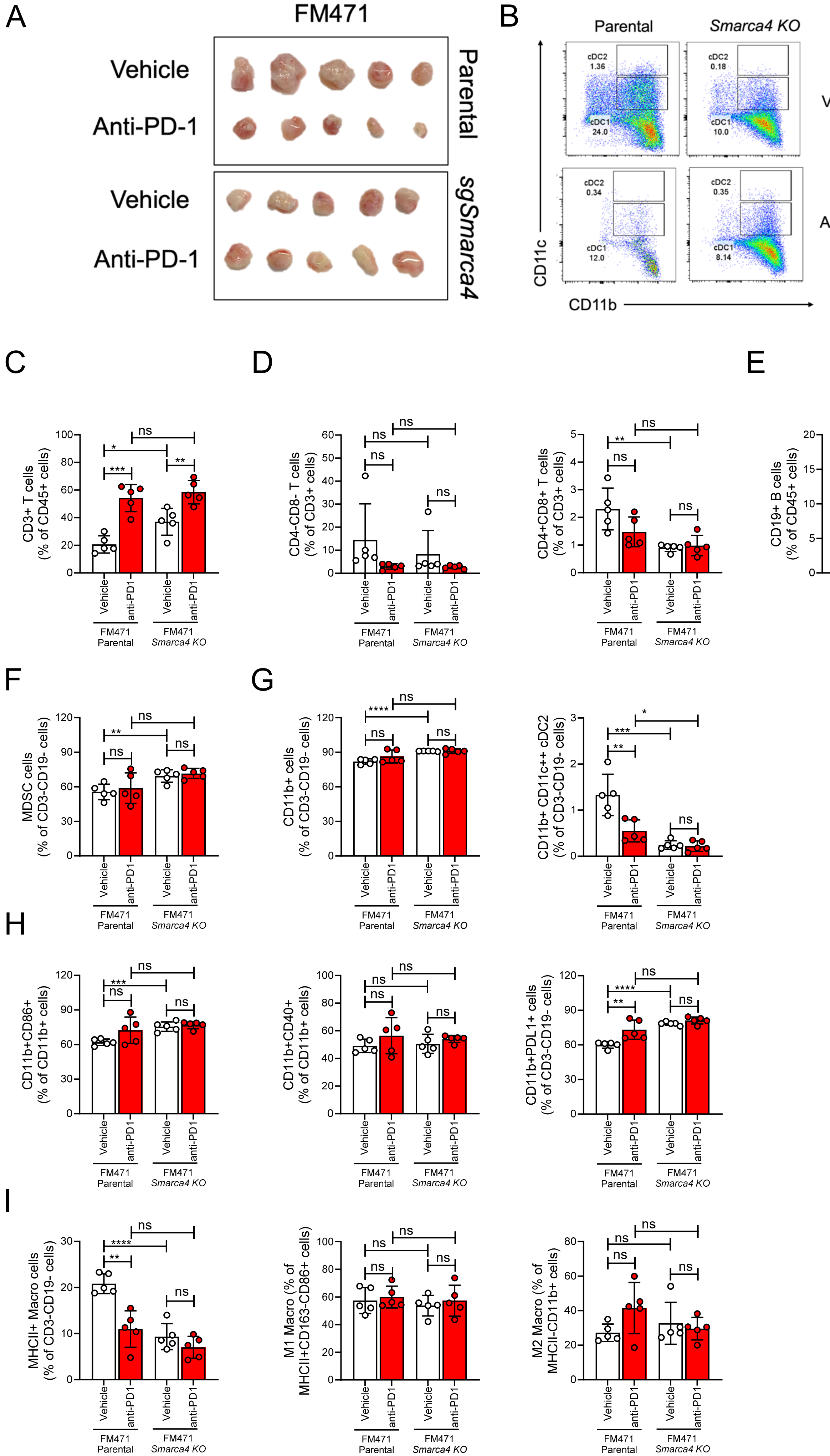


**Supplementary figure 3**

A

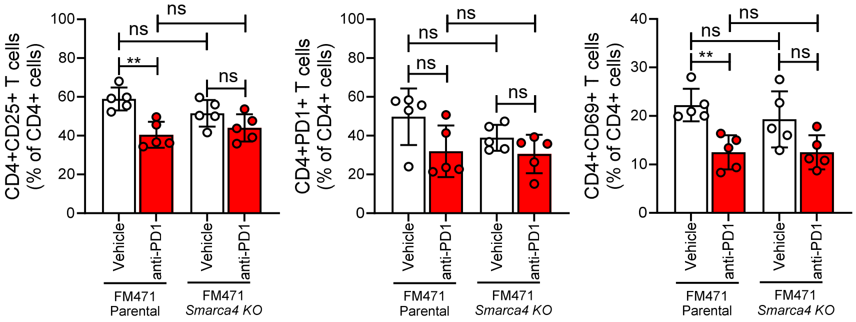


B

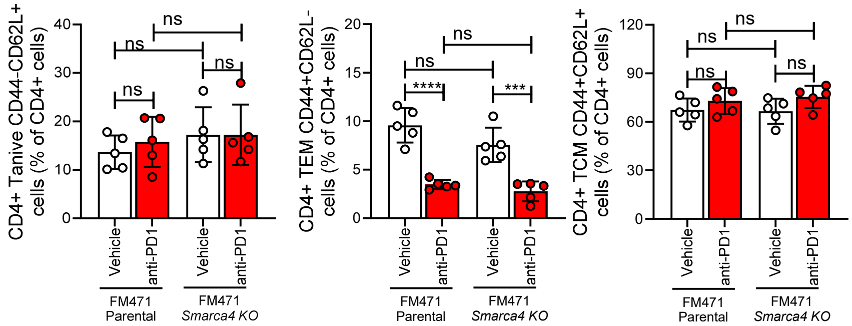


C

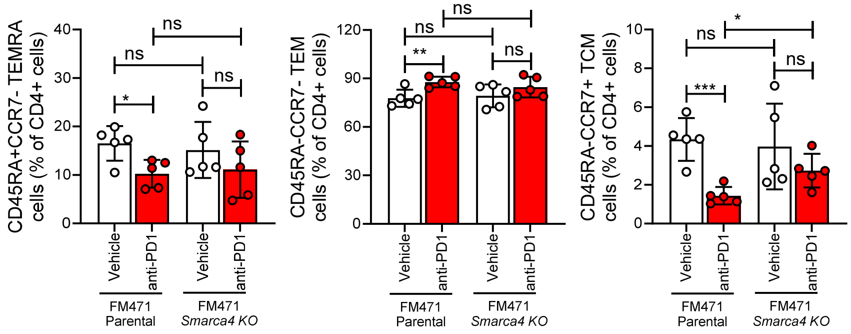


D

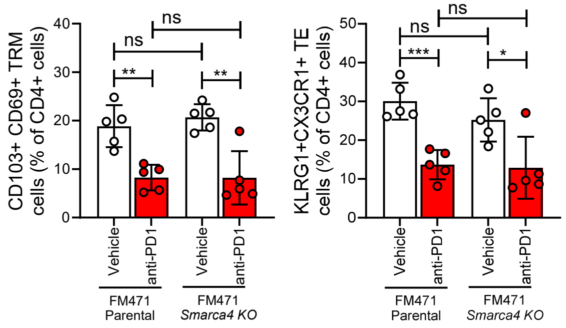


E

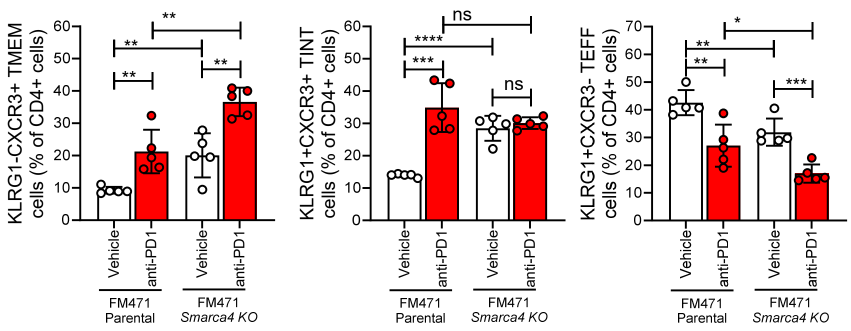


F

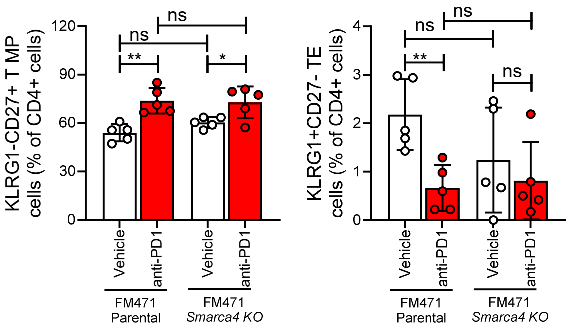


**Supplementary figure 4**

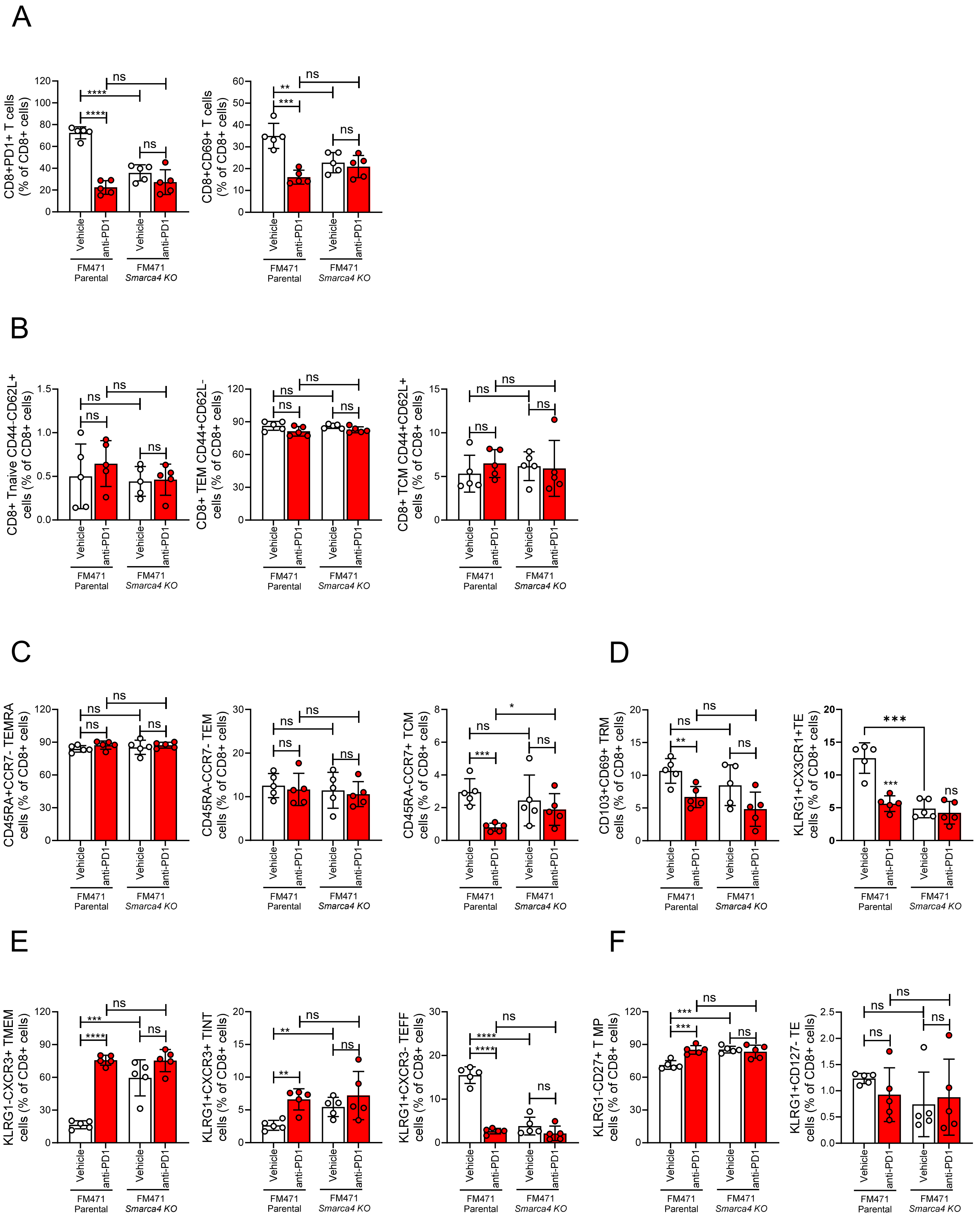


**Supplementary figure 5**

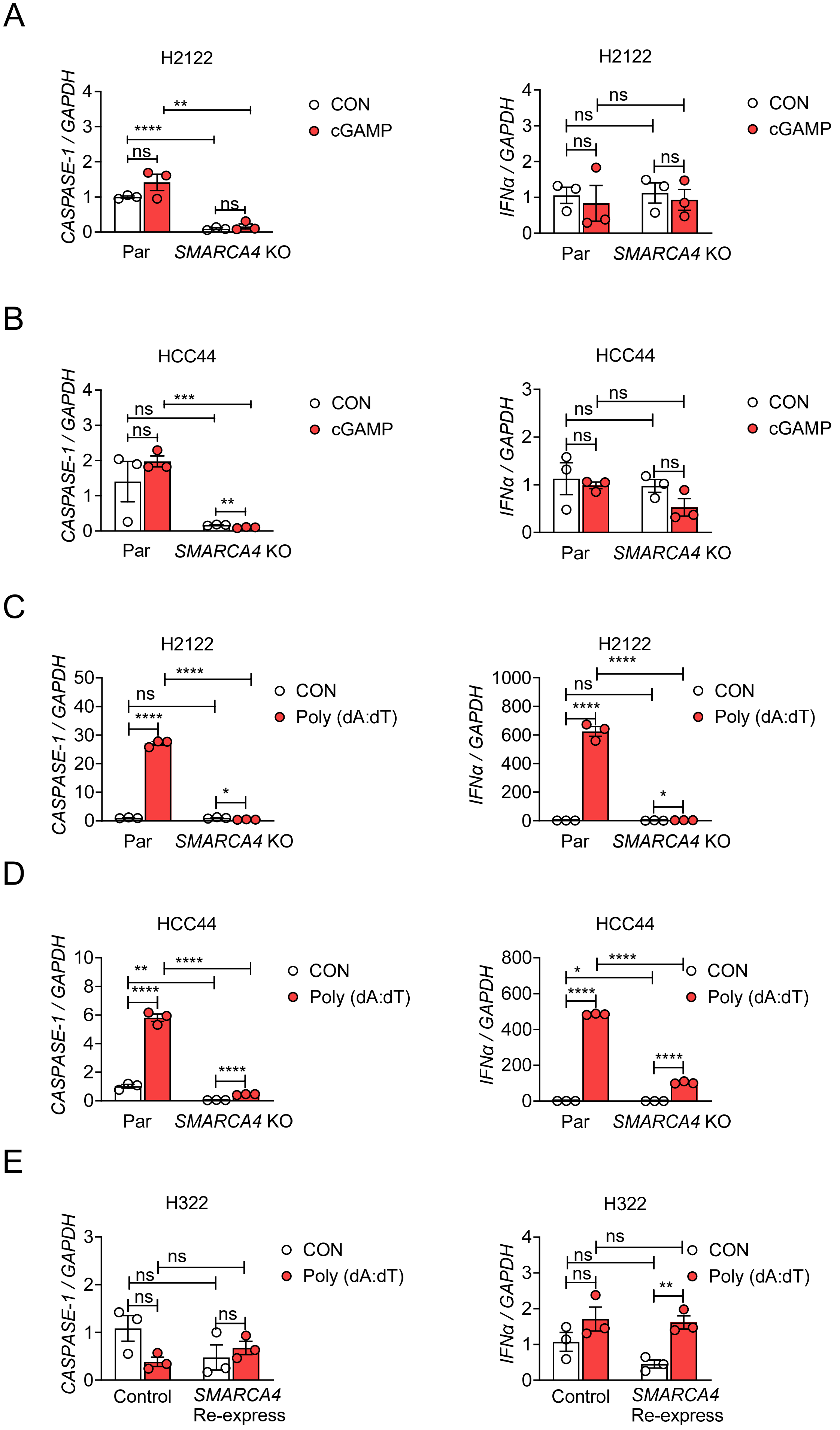


**Supplementary figure 6**
